# A humanized yeast model reveals dominant-negative properties of neuropathy-associated alanyl-tRNA synthetase mutations

**DOI:** 10.1101/2022.05.25.493316

**Authors:** Rebecca Meyer-Schuman, Sheila Marte, Tyler J. Smith, Shawna M.E. Feely, Marina Kennerson, Garth Nicholson, Mike E. Shy, Kristin S. Koutmou, Anthony Antonellis

## Abstract

Aminoacyl-tRNA synthetases (ARSs) are ubiquitously expressed, essential enzymes that ligate tRNA molecules to their cognate amino acids. Heterozygosity for missense variants or small in-frame deletions in five ARS genes causes axonal peripheral neuropathy, a disorder characterized by impaired neuronal function in the distal extremities. These variants reduce enzyme activity without significantly decreasing protein levels and reside in genes encoding homo-dimeric enzymes. These observations raise the possibility of a dominant-negative effect, in which non-functional mutant ARS subunits dimerize with wild-type ARS subunits and reduce overall ARS activity below 50%, breaching a threshold required for peripheral nerve axons. To test for these dominant-negative properties, we developed a humanized yeast assay to co-express pathogenic human alanyl-tRNA synthetase (*AARS1*) mutations with wild-type human *AARS1*. We show that multiple loss-of-function, pathogenic *AARS1* variants repress yeast growth in the presence of wild-type human *AARS1*. This growth defect is rescued when these variants are placed *in cis* with a mutation that reduces dimerization with the wild-type subunit, demonstrating that the interaction between mutant AARS1 and wild-type AARS1 is responsible for the repressed growth. This demonstrates that neuropathy-associated *AARS1* variants exert a dominant-negative effect, which supports a common, loss-of-function mechanism for ARS-mediated dominant peripheral neuropathy.

## INTRODUCTION

Hereditary peripheral neuropathies are a group of phenotypically and genetically heterogeneous diseases that are characterized by decreased sensory and/or motor axon function in the distal extremities. This leads to sensory loss and muscle atrophy, which often begins in the feet and lower legs, and may progress to include the hands and forearms of the upper extremities (1, 2). The most common type of inherited peripheral neuropathy is Charcot-Marie-Tooth (CMT) disease, which is estimated to affect between 1 in 1,200 and 1 in 2,500 individuals (3, 4).

In total, mutations in over 100 genes have been associated with CMT disease or related inherited peripheral neuropathies (5). Among these are the aminoacyl-tRNA synthetase (ARS) genes, a 37-member gene family that encodes ubiquitously expressed, essential enzymes. ARSs act as either monomers or oligomers to ligate tRNA to cognate amino acids, forming a critical substrate for protein translation (6). Variants in five aminoacyl-tRNA synthetases have been linked to dominant peripheral neuropathies: alanyl-(*AARS1*) (7), histidyl-(*HARS1*) (8), glycyl-(*GARS1*) (9), tryptophanyl-(*WARS1*) (10), and tyrosyl-(*YARS1*) tRNA synthetase (11). Although methionyl-tRNA synthetase (*MARS1*) variants have also been identified in patients with CMT disease (12–13), the genetic evidence for pathogenicity of these alleles is still incomplete. It remains to be seen how many additional ARSs will be implicated in dominant peripheral neuropathy. Further defining the locus and allelic heterogeneity of ARS-related neuropathy will be critical for patient diagnosis and defining disease mechanisms.

ARS activity is essential for cellular function. Indeed, bi-allelic ARS variants that reduce enzyme function cause early-onset recessive disorders that affect multiple tissues (14–15). In many cases, the constellation of recessive phenotypes includes peripheral neuropathy (16–18), demonstrating that peripheral neurons are sensitive to a reduction of ARS function. However, this reduction must be more than 50%, because heterozygosity for a null ARS allele is not sufficient to cause a penetrant neuropathic phenotype; mono-allelic null alleles are not found in CMT patient populations, but are found in unaffected individuals. Additionally, mice that are heterozygous for a *Gars1* null allele do not develop a peripheral neuropathy, while those carrying loss-of-function missense and in-frame deletion alleles do develop this phenotype (19–21). Based on these observations, haploinsufficiency is an unlikely disease mechanism for ARS-associated dominant neuropathy.

The pathogenic ARS variants linked to dominant peripheral neuropathy are exclusively missense mutations or small in-frame deletions (22). The absence of frameshift alleles and premature stop codons in patient populations indicates that an expressed mutant protein is required for pathogenicity. It is currently unknown if the dozens of identified pathogenic variants across the five implicated ARS genes (10, 21–28) exert a similar effect on a shared pathway, or if there are unique pathogenic mechanisms for each locus. One proposed mechanism is a neo-morphic gain-of-function effect, in which mutations expose novel protein interfaces that facilitate aberrant protein-protein interactions, leading to dysregulated neuronal pathways (29–33). However, it is unlikely that the dozens of different alleles across all five ARS loci give rise to an identical gain-of-function effect via aberrant interactions. Rather, any common mechanism would likely be related to the shared canonical function of charging tRNA in the cytoplasm and axoplasm of peripheral neurons.

One notable commonality is that all five enzymes function as homodimers. This raises the possibility of a dominant-negative mechanism, in which decreased function of the mutant subunit reduces the function of the wild-type subunit in the dimer; this would lower the overall ARS activity in the cell below 50%. This mechanism is supported by an abundance of data showing that the majority of neuropathy-associated ARS variants reduce gene or enzyme function (10, 21–23, 25–27). Additionally, pathogenic variants do not decrease protein levels (20, 21, 34, 35) nor do they abolish dimerization (10, 11, 27, 36, 37). It has also been shown that pathogenic ARS variants impair protein synthesis in affected neurons. Expressing neuropathy-associated ARS alleles decreases global protein translation *in vitro* (10, 36, 38) and *in vivo* (39, 40), increases eIF2-alpha phosphorylation (36, 38, 40, 41) and activates the integrated stress response (38, 40, 41). Importantly, these phenotypes can be recapitulated *in vitro* by chemical inhibition of the affected ARS enzyme (36), which directly links these phenotypes to reduced enzyme function. For a subset of *GARS1* mutations, the loss of function has been defined as altered affinity for tRNA^Gly^ (38, 41), which depletes the cell of charged tRNA^Gly^ and causes ribosome stalling at glycine codons (38, 41). Cumulatively, these studies provide compelling evidence for a loss-of-function mechanism. Considering the evidence against haploinsufficiency, it is possible that a loss-of-function effect is mediated by pathogenic ARS alleles interfering with the function of the wild-type allele in a dominant-negative fashion. In support of this, previous studies in yeast have shown that cells co-expressing both wild-type and mutant yeast tyrosyl-tRNA synthetase show impaired growth, compared to cells expressing only wild-type tyrosyl-tRNA synthetase (11). However, it remains unclear if these phenotypes are due to a dominant-negative effect or an unrelated toxic effect of the mutant allele. Addressing this question will attempt to identify a unifying mechanism of disease for all five implicated dimeric ARS enzymes. Furthermore, developing an approach to systematically assess ARS variants for a dominant-negative effect will provide a relevant framework to assess the pathogenicity of newly identified ARS variants in patients with dominant CMT disease.

Here, we report a yeast model to test human *AARS1* variants for a dominant-negative effect. We focused on well-characterized alleles in the anti-codon binding domain (R329H [7, 22, 42, 43]) and the amino acid activation domain (G102R [44]). We found that R329H and G102R, as well as three additional *AARS1* variants (R326W, R329S, and R329C) have a dominant, repressive effect on yeast cell growth when co-expressed with wild-type human *AARS1*. We then engineered a dimer-disrupting variant in the C-terminal domain and modelled it in *cis* with the pathogenic *AARS1* variants. These double-mutants rescued the impaired yeast growth, demonstrating that the dominant effect of mutant AARS1 is dependent on dimerization with wild-type AARS1 and that neuropathy-associated *AARS1* variants can be classified as dominant-negative (or antimorphic) alleles.

## MATERIAL AND METHODS

### Yeast vector construction and growth assays

All yeast assays were performed using the ptetO7-*ALA1* strain from the Yeast Tet-Promoters Hughes Collection (YSC1180-202219317, Horizon Discovery). *AARS1* variants were generated using site-directed mutagenesis (Agilent QuikChange II XL Site-Directed Mutagenesis Kit) against the *AARS1* open reading frame in pDONR221 (primers available upon request). All clones were verified via Sanger sequencing. The Gateway cloning (Invitrogen) LR reaction was used to recombine the wildtype or mutant *AARS1* locus into pAG425GAL-ccdB (Addgene #14153). This is a Gateway-compatible vector with a 2-micron origin of replication that produces a high vector copy number per cell, a *GAL1* promoter that drives high expression of the target gene in a galactose-inducible fashion, and a *LEU2* auxotrophic marker.

To assess the function of *AARS1* variants independent of wild-type *AARS1*, a p413 vector (ATCC #87370) with no *AARS1* insert (‘Empty’) was introduced into the ptetO7-*ALA1* strain using lithium acetate yeast transformation. The p413 vector contains an *ADH1* promoter to drive constitutive expression of the target gene, a centromeric origin of replication to produce a low plasmid copy number per cell, and a *HIS3* auxotrophic marker for selection. This transformation was followed by a second transformation to introduce a pAG425 vector harboring wild-type or mutant *AARS1* (see above). Colonies were grown on media lacking histidine and leucine (DO Supplement -His/-Leu, Takara Bio) to select for the presence of both vectors. After transformation, colonies were grown in 2mL liquid media in a 14mL round-bottom conical tube (Fisher Scientific) for two days at 30°C, shaking at 275 rpm (on an I-24 incubator shaker, New Brunswick Scientific) until saturated. Yeast were then diluted 1:10, 1:100, and 1:1000 in water. 10μl of undiluted or diluted samples were spotted on plates containing glucose, galactose/raffinose (Takara Bio Minimal SD Bases), or galactose/raffinose with 10μg/ml doxycycline (Fisher Scientific BP26531). Plates were imaged after four days of growth.

To co-express mutant and wild-type *AARS1* in yeast, Gateway Cassette C (Invitrogen) was cloned into the p413 vector just downstream of (3’ to) the *ADH1* promoter using the *Sma*I restriction site. Clones were sequence verified to confirm correct orientation. The LR Gateway reaction was then used to recombine wild-type *AARS1* from pDONR221 into p413. This construct was transformed into the ptetO7-ALA1 strain, followed by *AARS1* (wild-type or mutant) in pAG425. Yeast were grown and spotted as detailed above. All growth assays (studying a single *AARS1* allele or two co-expressed *AARS1* alleles) were performed side-by-side with the same pAG425 plasmid aliquots to enable direct comparisons. To quantitatively analyze yeast spots, mean gray values of each spot were obtained using ImageJ, according to previously published protocol (45).

### Yeast protein isolation

To assess the expression of each human AARS1 protein in yeast, the ptetO7-*ALA1* strain was transformed with a vector (pAG425) to express wild-type or mutant *AARS1* and grown on media lacking leucine (DO Supplement -Leu, Takara Bio). One colony was picked, placed into 3mL media, and grown for 2-3 days shaking at 275 rpm at 30°C until saturated, reaching an optical density (OD_600_) of approximately 2. Yeast were then centrifuged at 1,000xg for 10min, washed once with water, transferred to a 1.5mL Eppendorf tube, then centrifuged at 21,130xg for 1min. The supernatant was removed and the pellet was stored at −80°C. The pellet was thawed in 150μl yeast lysis buffer (50mM Na-HEPES pH 7.5, 100mM NaOAc, 1mM EDTA, 1mM EGTA, 5 mM MgOAc, 5% glycerol, 0.25% NP-40, 3 mM DTT) with 1X Halt Protease Inhibitor Cocktail (Thermo Fisher Scientific). Approximately 100μl of 0.5mm cold glass beads (Biospec Products) were added to each sample. Samples were vortexed at 4°C for 3min, followed by 2min resting on ice, followed by 3min of vortexing at 4°C. To remove the lysate from the beads, a 26-gauge needle (BD PrecisionGlide) was used to make a hole in the bottom of the 1.5mL tube, which was then immediately inserted into a 14mL round bottom conical tube. Lysates were centrifuged at 200xg at 4°C for 5min. The lysates were collected from the bottom of the conical tube and transferred to a 1.5mL Eppendorf tube, and were then centrifuged at 16, 363xg for 5min at 4°C. Supernatants were collected and protein concentrations were measured using the Thermo Scientific Pierce BCA Protein Assay kit, and 50μg of protein per sample was used for western blot analysis (see below).

### Co-immunoprecipitation of wild-type AARS1 and mutant AARS1

To test for interactions between wild-type and mutant AARS1 proteins we performed coimmunoprecipitation followed by western blot analysis in cultured mammalian cells. The LR Gateway reaction was used to recombine the wild-type or mutant *AARS1* open reading frame from pDONR221 into pDEST40 (Thermo Fisher Scientific) or pTM3xFLAG (gift from Moran Laboratory, University of Michigan). These vectors allow differential tagging of the mutant and wild-type *AARS1* alleles; wildtype AARS1-3xFLAG (a C-terminal tag) was expressed from pTM3xFLAG using a CMV promoter, and either wild-type or mutant AARS1-6xHis (a C-terminal tag) was expressed from pDEST40 using a CMV promoter. 100mm plates (Falcon) were seeded with 1.5-2 million HEK293T cells; the following day, these were transfected with 0.5pmol plasmid using Lipofectamine 3000 (Invitrogen). 48h after transfection, cells were harvested using Trypsin-EDTA (Gibco, Fisher Scientific) and centrifuged at 800xg for 2min at 4°C. Cells were then washed once with 1X PBS (Thermo Fisher Scientific), centrifuged again (as above), and then resuspended in 1mL lysis buffer (20mM Tris-HCl pH 8, 137mM NaCl, 2mM EDTA, 1% NP-40, 0.25% sodium deoxycholate) with 1X Halt Protease Inhibitor Cocktail (Thermo Fisher Scientific). Samples were incubated for 2h, rocking at 4°C, then centrifuged for 15min at 16,363xg at 4°C. The supernatant was collected and protein concentration was measured using the Thermo Scientific Pierce BCA Protein Assay kit.

To conjugate beads with individual antibodies, 25μl of Dynabeads Protein G (Fisher Scientific) were aliquoted per sample. All immunoprecipitations were performed using a MagnaRack (Invitrogen). Each aliquot was washed twice with 500μl conjugation buffer (0.5% BSA, 0.1% Triton X-100 in PBS), then suspended in 500μl conjugation buffer with 2μg 6xHis antibody (Abcam 18184) or 2μg FLAG antibody (BioLegend 637302). Beads and antibody were incubated overnight with rocking at 4°C.

Prior to immunoprecipitation, lysates were pre-cleared to remove any proteins with non-specific affinity for the magnetic beads. An additional 25μl of Dynabeads per sample was aliquoted and washed once with lysis buffer. Then, 1mg of cell lysate in 500μl lysis buffer was added to the beads and rocked at 4°C for 2h. Supernatant from the antibody-conjugated beads was then removed, and the pre-cleared lysates were added. Samples were incubated for 3h rocking at 4°C. For anti-6xHis IPs, samples were washed four times with 1mL high salt buffer (10mM Tris-HCl pH 7.5, 400mM NaCl, 1 mM EDTA, 1mM EGTA, 0.5% NP-40). For anti-FLAG IPs, samples were washed three times with 1mL low salt buffer (10mM Tris-HCl pH 7.5, 137mM NaCl, 1 mM EDTA, 1mM EGTA, 0.5% NP-40). On the last wash, samples were moved to fresh 1.5 mL tubes to prevent co-elution of proteins bound to the tube walls. Samples were re-suspended in 50μl wash buffer with 50μl 2x Tris Glycine Buffer (Invitrogen). 4μl BME was added before samples were boiled at 99°C for 5min and the supernatant was collected for western blot (see below). Samples were divided in half and loaded in duplicate for immunoblotting with anti-AARS1, anti-6xHis, or anti-FLAG.

### Disuccinimidyl suberate crosslinking

To determine the degree of AARS1 dimerization in patient cells, AARS1 protein was crosslinked with disuccinimidyl suberate (DSS) and analyzed by western blot. Patient and control fibroblasts were grown at 37°C in 5% CO_2_ and standard growth media (DMEM supplemented with 10% FBS, 2mM L-glutamine, 100U/mL penicillin, and 50μg/mL streptomycin [Invitrogen]). Approximately 1 million cells were harvested from each sample with Trypsin-EDTA (Gibco, Fisher Scientific) and centrifuged at 800xg for 2min at 4°C. They were then washed once with 1X PBS (Thermo Fisher Scientific), transferred to a 1.5mL tube, and centrifuged again (as above). Cells were then re-suspended in 50mM HEPES 0.5% NP-40. The sample was divided in two and 50mM DSS (Thermo Fisher Scientific) was added to one aliquot to a final concentration of 5mM. Both aliquots were incubated at room temperature for 30min. The crosslinking reaction was then quenched with a final concentration of 30mM TrisCl pH 7.5 at room temperature for 15min. Samples were centrifuged at 16,363xg for 10 min at 4°C, and the supernatant was collected for western blot analysis (see below). 20μg of protein was analyzed for each sample. An antibody against human AARS1 (Bethyl Laboratories A303-473A) was used at a dilution of 1:500.

### Co-immunoprecipitation of wild-type ALA1 and wild-type AARS1

To investigate an interaction between yeast ALA1 and human AARS1, co-immunoprecipitation experiments were performed. First, the endogenous yeast *ALA1* coding sequence was amplified from a previously published (42) pDONR221 clone with or without a C-terminal 6xHis tag encoded in the reverse primer (primer sequences available upon request). Then, Gateway cloning was used to recombine these constructs into p413 (see above). The ptetO7-*ALA1* strain was transformed with p413 to express either 6xHis-tagged or untagged *ALA1*, then subsequently transformed with pAG425 to express either R329H or G757* human *AARS1*. Colonies were grown for 2-3 days until saturated in -leu -his liquid glucose growth medium, then washed in water and re-suspended in 125-250mL -leu - his galactose liquid culture (Takara Bio Minimal SD Bases, Takara Bio DO Supplement -His/-Leu). Cultures were grown to saturation, then centrifuged at 1000xg at 4°C for 20min. Yeast were washed with water and aliquoted evenly into 4-5 1.5mL tubes. Each sample was then centrifuged at 21,130xg for 1min. The supernatant was removed and pellets were stored at −80°C. The pellets were thawed in yeast lysis buffer (50mM Na-HEPES pH 7.5, 100mM NaOAc, 1mM EDTA, 1mM EGTA, 5 mM MgOAc, 5% glycerol, 0.25% NP-40, 3 mM DTT) with 1X Halt Protease Inhibitor Cocktail (Thermo Fisher Scientific). Approximately 100μl of buffer was used for each 100mg of pellet. Cells were lysed using the methods detailed above.

25μl of Dynabeads Protein G (Fisher Scientific) were prepared for each of the samples. Beads were washed twice with 500μl conjugation buffer (0.5% BSA, 0.1% Triton X-100 in PBS), then resuspended in 500μl buffer and 2μg anti-AARS1 (ab226259). Beads and antibody were incubated overnight with rocking at 4°C. Yeast cell lysates were pre-cleared before immunoprecipitation: 25μl of magnetic beads were aliquoted and washed once with lysis buffer, before a 2mg aliquot of yeast lysate in a total of 500μl lysis buffer was added. Samples were incubated with rocking at 4°C for 1h. The supernatant was then removed from antibody-conjugated beads and replaced with the precleared lysates. These samples were then incubated with rocking for 2.5h at 4°C. After incubation, samples were washed once with 500μl lysis buffer, once with 200μl lysis buffer, and then resuspended in 100μl lysis buffer before being transferred to a fresh 1.5mL Eppendorf tube. The supernatant was then removed and beads were suspended in 25μl lysis buffer and 25μl 2x Tris Glycine Buffer (Invitrogen). Samples were boiled for 5min with 2μl BME and supernatants were removed to analyze in western blot assays.

### Western blot analyses

To assess the levels of specific proteins in each experiment, western blots were performed. Protein concentrations for each sample were measured using the Thermo Scientific Pierce BCA Protein Assay kit. Samples were prepared with 1X Novex Tris-Glycine SDS sample buffer (Invitrogen) and 2-mercaptoethanol (BME), and boiled at 99°C for 5min. Protein samples were separated on precast 4-20% Novex Wedgewell Tris-glycine gels (Invitrogen) at 150V for 1h and 15min. PVDF membranes (Millipore Sigma) were pre-washed in 100% methanol for 1min, then soaked in 1X transfer buffer (Invitrogen) and 10% methanol between two pieces of filter paper (Thermo Fisher Scientific). The separated protein samples were transferred to the PVDF membranes using a Mini Trans-Blot Electrophoretic Transfer Cell (Biorad) at 100V for 1h. Membranes were then blocked for 1h with a 5% milk powder solution in 1X TBST. Primary antibodies were applied in 5% milk and membranes were incubated by rocking overnight at 4°C. The following day, membranes were washed three times with 1X TBST. Secondary antibodies (Licor) against mouse (for the 6xHis primary antibody and the PGK1 primary antibody), rabbit (for the AARS1 primary antibody and the actin primary antibody), or rat (for the FLAG primary antibody) were diluted in 5% milk powder solution at a concentration of 1:20,000, along with 0.1% Tween-20 and 0.02% SDS. This solution was applied to membranes for 1h, rocking at room temperature. Membranes were then washed three times with 1X TBST before exposure using a Licor Odyssey CLx Imaging System.

For yeast protein and HEK293T co-immunoprecipitation experiments, the AARS1 antibody (Bethyl Laboratories A303-473A) was used at 1:1,000 dilution. For fibroblast DSS assays, the same AARS1 antibody was used at 1:500 dilution. For HEK293T and yeast co-immunoprecipitation experiments, the 6xHis antibody (Abcam 18184) was used at a dilution of 1:3,000. The FLAG antibody (BioLegend 637302) was used at a 1:2,500 dilution. The loading control was actin (Sigma A5060, 1:5,000) for mammalian protein blots and PGK1 (Abcam ab113687, 1:3,000) for yeast protein blots. For coimmunoprecipitation studies of AARS1 and ALA1, the AARS1 antibody used was Abcam ab226259 at a dilution of 1:500.

### AARS1 protein structure prediction

To predict whether Q855* significantly alters AARS1 protein structure, the AlphaFold Colab notebook (46) was used to predict the protein structure of the first 854 amino acids of AARS1 (Uniprot P49588-1). To enable a direct comparison between mutant and wild-type, AlphaFold Colab was also used to predict the structure of the full-length AARS1 amino acid sequence. Predicted structures were visualized and aligned using PyMOL.

## RESULTS

### Pathogenic *AARS1* alleles repress yeast cell growth in the presence of wild-type *AARS1*

Baker’s yeast (*Saccharomyces cerevisiae*) has proven to be an effective model to study the functional consequences of pathogenic ARS alleles in a living cell (22). To investigate dominant-negative properties of neuropathy-associated *AARS1* variants, we developed an assay to test the effects of coexpressing human wild-type *AARS1* and human mutant *AARS1* on yeast viability, using the ptetO7-*ALA1* strain. In this strain, the yeast *AARS1* ortholog, *ALA1*, is placed under control of a doxycycline-repressible promoter (47). We transformed this strain with: (**1**) a low-copy, centromere-bearing vector (p413 [48]) containing a wild-type *AARS1* allele; and (**2**) a high-copy number vector (*i.e*., bearing a 2 micron origin of replication) with a galactose-inducible promoter (pAG425 [49]) directing high levels of expression of either wild-type or mutant *AARS1* alleles (Supplemental Figure 1B). To test mutant *AARS1* alleles for a dominant effect on yeast cell growth, yeast cells were grown in the presence of galactose (to express wild-type or mutant *AARS1* from the pAG425 vector) and doxycycline (to repress endogenous *ALA1*); in this system, wild-type *AARS1* is constitutively expressed from p413. Subsequent yeast growth was then solely dependent on the two forms of human *AARS1:* one wildtype (from p413) and one wild-type or mutant (from pAG425).

**Figure 1.**
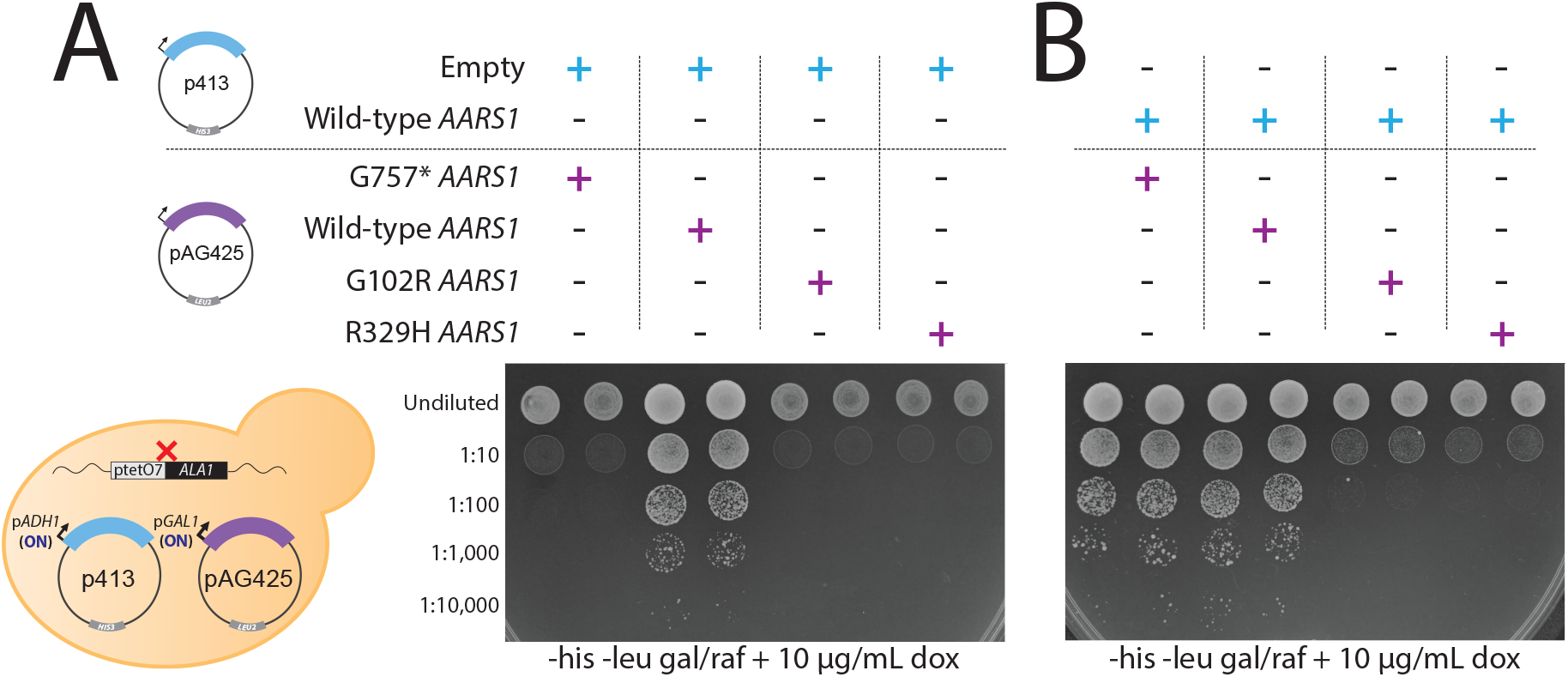
G102R and R329H *AARS1* are loss-of-function alleles that repress yeast cell growth in the presence of wild-type *AARS1*. (**A**) Yeast harboring an endogenous doxycycline-repressible *ALA1* locus were transformed with a p413 vector with no insert and a pAG425 vector to express either wildtype or mutant human *AARS1*. Cultures were plated undiluted or diluted on media lacking histidine and leucine, and containing galactose/raffinose and doxycycline. (**B**) Similar experiment as shown in panel A; here, yeast were first transformed with a p413 vector expressing wild-type human AARS1. For both panels, the vectors present in each experiment are indicated across the top, the dilution of the spotted yeast cultured is indicated on the left side of the image, and the media conditions are indicated across the bottom of the image (his = histidine; leu = leucine; gal = galactose; raf = raffinose; dox = doxycycline). Representative images are shown from thirteen (for G102R) or sixteen (for R329H) biological replicates. A cartoon on the bottom left illustrates the experimental conditions for all samples.

To evaluate neuropathy-associated *AARS1* variants for a dominant-negative effect, we focused on two well-characterized pathogenic *AARS1* variants, R329H and G102R. R329H is a recurrent mutation in the tRNA recognition domain that has been identified in 10 families with dominant, axonal CMT (7, 24, 42, 43, 50). It affects a highly conserved residue and significantly impairs AARS1 enzymatic function when assessed via *in vitro* aminoacylation assays under Michaelis-Menten conditions (42). The G102R *AARS1* variant affects a highly conserved residue in the activation domain of AARS1 and was found in a single pedigree with dominant myeloneuropathy (44). Both G102R and R329H have been modelled in the yeast ortholog *ALA1*, and were unable to support yeast growth, indicating a loss-of-function effect *in vivo* (42, 44) To distinguish dominant-negative properties of these variants from a strictly loss-of-function effect, we generated a premature stop codon, G757* (Supplemental Figure 1A), which does not generate detectable levels of AARS1 protein (Supplemental Figure 1B). Therefore, this variant is expected to be a loss-of-function allele that does not exert dominant-negative effects. This variant is a more precise negative control than an empty vector because it includes the *AARS1* coding sequence. Yeast transformed with the G757* allele must replicate a nearly identical DNA sequence, and express a nearly identical mRNA sequence, as yeast transformed with the G102R or R329H alleles.

To confirm that R329H and G102R human *AARS1* are loss-of-function alleles in a yeast complementation assay, ptetO7-*ALA1* yeast were first transformed with an empty p413 vector, then with pAG425 expressing either wild-type or mutant human *AARS1*. When these strains were plated on media containing glucose, there were no observable growth defects (Supplemental Figure 2A). When these strains were plated on media containing galactose and doxycycline, the yeast expressing G757* did not display any visible growth (Figure 1A), indicating that yeast cannot grow without *ALA1* expression (*i.e*., without functional AARS1 protein). Transformation with wild-type *AARS1* lead to robust yeast growth, confirming that wild-type human *AARS1* can complement loss of yeast *ALA1*. Neither G102R nor R329H *AARS1* supported yeast growth (Figure 1A), confirming previous reports that these are loss-of-function alleles *in vivo* (42, 44)

To determine if mutant *AARS1* alleles can repress the function of wild-type *AARS1*, ptetO7-*ALA1* yeast cells were transformed with wild-type *AARS1* on the low-copy p413 vector, then with wild-type or mutant *AARS1* on the high-copy pAG425 vector. Transformed strains spotted on glucose media displayed no observable growth defects (Supplemental Figure 2B). These strains were then spotted on galactose (to induce expression from the pAG425 vector) and doxycycline (to repress *ALA1* expression). Co-expression of wild-type and G757* *AARS1* supported robust growth, as did coexpression of the two wild-type *AARS1* plasmids (Figure 1B). These data indicate that the G757* null allele has no negative impact on yeast cell growth and, similarly, that expressing two copies of wildtype *AARS1* does not repress yeast growth. In contrast, co-expression of wild-type *AARS1* with either G102R or R329H *AARS1* caused significantly reduced yeast growth (Figure 1B). These data demonstrate that the two pathogenic, neuropathy-associated alleles are not only loss-of-function alleles but that they are also detrimental to yeast cell growth in the presence of wild-type *AARS1*. These observations are consistent with mutant AARS1 interfering with the function of the wild-type AARS1 protein. Importantly, the mutation-associated growth deficit is partially rescued by restoring *ALA1* expression (Supplemental Figure 3A), which increases growth 23-fold for yeast expressing G102R and 11-fold for yeast expressing R329H (Supplemental Figure 3B). These data suggest that the dominant growth deficit caused by R329H and G102R *AARS1* is due to reduced alanine-tRNA charging. Interestingly, *ALA1* only partially rescues the growth deficit associated with R329H. This may be related to the observation that human R329H AARS1 interacts with ALA1 (Supplemental Figure 4B), suggesting that human R329H AARS1 has a dominant-negative effect on wild-type yeast ALA1.

### Pathogenic *AARS1* variants do not significantly reduce dimerization

The data presented above demonstrate that the pathogenic, loss-of-function *AARS1* variants G102R and R329H *AARS1* dominantly repress yeast cell growth in the presence of wild-type *AARS1*, consistent with a dominant-negative effect. However, the data do not rule out some other form of gain-of-function toxicity unrelated to AARS1 function. To directly test for a dominant-negative effect, we first investigated if mutant AARS1 dimerizes with wild-type AARS1. Although ultracentrifugation analyses have demonstrated that isolated mutant ARS proteins retain homo-dimerization (27, 36, 37), no studies have addressed hetero-dimerization between the AARS1 mutant and wild-type subunits. To address this, HEK293T cells were transfected with a vector expressing wild-type human AARS1 with an in-frame C-terminal 3xFLAG tag and with a vector expressing mutant AARS1 (G102R or R329H) with an in-frame C-terminal 6xHis tag (Figure 2A). After growth for 48h, cells were lysed and AARS1-6xHis was immunoprecipitated. Co-immunoprecipitated proteins were subjected to western blot analysis with an anti-FLAG antibody to detect AARS1-3xFLAG. The reciprocal immunoprecipitation was also performed by immunoprecipitating AARS-3xFLAG and immunoblotting for AARS-6xHis. Both approaches detected comparable interactions between wild-type and wild-type, wild-type and G102R, and wild-type and R329H (Figure 2B and 2C). To assess dimerization of endogenous AARS1 in patient cells, fibroblasts from a patient heterozygous for R329H *AARS1* were crosslinked with disuccinimidyl suberate (DSS), along with two independent control fibroblast cell lines. In untreated lysates (-DSS), AARS1 protein is detected between 100 and 130 kDa, consistent with its predicted size of 107 kDa. In DSS-treated lysates, an additional band migrates at approximately twice the molecular weight, consistent with a dimeric AARS1 protein (Figure 2D). The percentage of AARS1 in dimeric form was not significantly different between control and patient cell lines (Figure 2E). Combined, these data indicate that mutant AARS1 proteins retain the ability to dimerize with wild-type AARS1, which is required for a dominant-negative effect.

**Figure 2.**
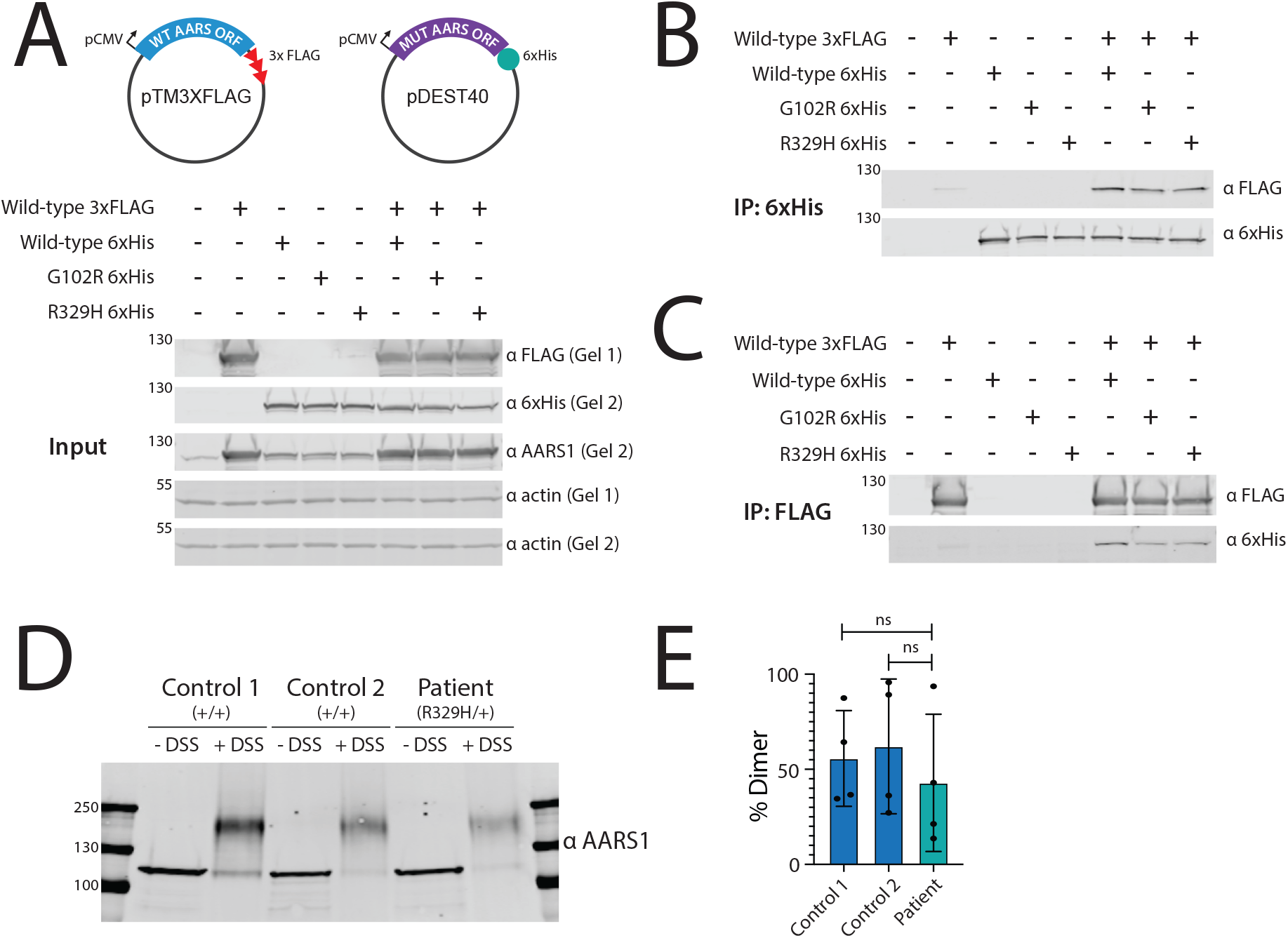
G102R and R329H AARS1 dimerize with wild-type AARS1. **(A)** HEK293T cells were transfected with vectors to co-express wild-type and mutant human *AARS1*, and a western blot was performed to detect the resulting proteins along with endogenous loading controls. The image is representative of three independent replicates. A cartoon along the top illustrates the constructs employed in the experiments, and the presence or absence of each construct is indicated across the top of the gel image. Protein molecular weights are indicated in kilodaltons (kDa) along the left side of the image and antibodies are indicated along the right side. (**B**) After immunoprecipitation with an anti-6xHis antibody, a western blot was performed to detect co-immunoprecipitated proteins. A representative image from five (for R329H) or three (for G102R) independent replicates are shown. This image is annotated as in panel A. (**C**) After immunoprecipitation with an anti-FLAG antibody, a western blot was performed to detect co-immunoprecipitated proteins. A representative image from two independent replicates is shown. This image is annotated as in panels A and B. **(D)** After treating patient and control samples with a protein cross-linking agent, a western blot was performed to detect endogenous AARS1 protein. The image is representative of four independent technical replicates. Samples and conditions are indicated across the top of the image, protein molecular weights are indicated in kilodaltons (kDa) along the left side, and the antibody employed is indicated on the right side. DSS = disuccinimidyl suberate. **(E)** The percentage of dimeric AARS1 protein signal in the total AARS1 protein signal (panel D) was quantified with ImageJ. The mean and standard deviation of four independent technical replicates is shown. Unpaired t-tests with Welch’s correction were performed to determine if there was a statistically significant difference between R329H/+ cells and either of the two controls. ns = not significant.

### Engineering dimer-disrupting *AARS1* variants

If neuropathy-associated *AARS1* variants (e.g. R329H or G102R) act in a dominant-negative manner, then introducing a dimer-disrupting variant in *cis* with the pathogenic variant should reduce interactions between mutant and wild-type AARS1 proteins, temper any dominant-negative effects, and rescue the observed reduction in yeast cell growth. To identify dimer-disrupting variants, a series of deletions were designed in the C-terminal domain of AARS1 based on the published crystal structure (51). The C-terminal domain is predicted to be essential for AARS1 dimerization, although it has been shown to be dispensable for AARS1 expression and tRNA charging activity *in vitro* (51). Engineered deletions in this domain targeted amino acids that have multiple contacts with the opposite subunit (Figure 3A, left). This series comprised a seven amino acid deletion to encompass several contact points (K943_Q949del) as well as smaller deletions within or near this region (N944_G946del and Q949_E950del). This series also included the deletion of a single codon, which encodes C947, a cysteine residue that forms a putative disulfide bond with C773 on the opposite subunit (51). Finally, a stop codon at Q855 was introduced to ablate the 113 amino-acid residues of the terminal globular domain (Figure 3A, right).

**Figure 3.**
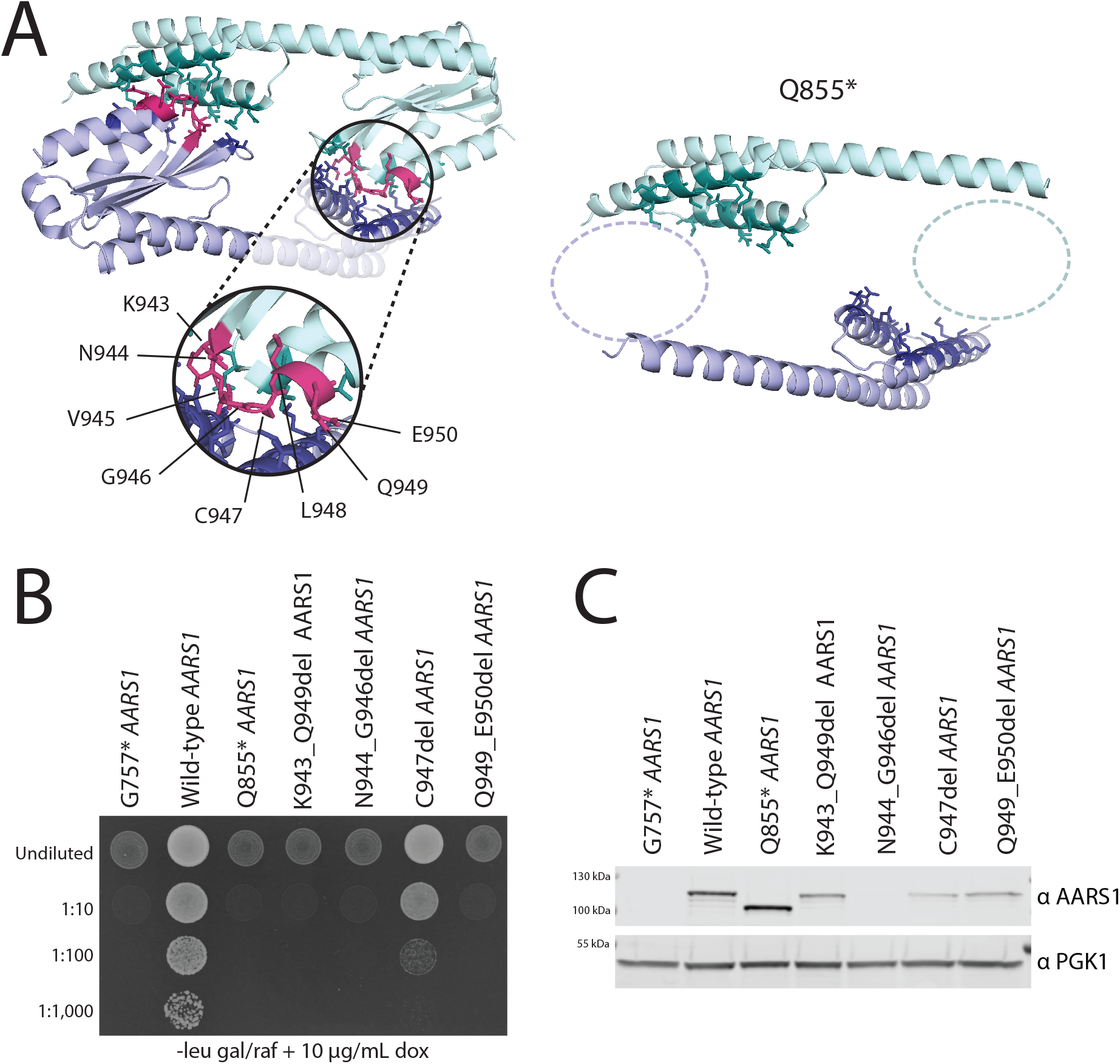
Engineering dimer-reducing *AARS1* variants. **(A)** A cartoon generated from PyMOL illustrates the crystal structure of the AARS1 C-terminal dimerization domain (left side). One subunit from the dimer is shown in green, the other in purple. Amino-acid residues that contact the opposite subunit are shown in dark green or dark purple. The residues targeted in this assay are shown in pink and labelled in the inset. On the right side of this panel is a second cartoon that illustrates the AARS1 C-terminal dimerization domain with the Q855* mutation. The dashed circles indicate the globular domain that is ablated by the premature stop codon. **(B)** Yeast harboring a doxycycline-repressible endogenous *ALA1* locus were transformed with a pAG425 vector to express either wild-type or mutant human *AARS1* (*i.e*., one of the engineered mutations affecting the residues highlighted in panel A). Cultures were plated undiluted or diluted on media lacking leucine, and containing galactose/raffinose and doxycycline. A representative image of four biological replicates is shown. The dilution of the spotted yeast cultured is indicated on the left and the media conditions are indicated across the bottom (leu = leucine; gal = galactose; raf = raffinose; dox = doxycycline). **(C)** Yeast protein lysates were subjected to western blot analysis to detect the human AARS1 proteins expressed from wild-type and mutant expression constructs, which are indicated across the top. Yeast were grown in galactose and raffinose media lacking leucine, with no doxycycline. A representative image of three biological replicates is shown.

To assess the ability of each of the five putative dimer-disrupting *AARS1* alleles to support yeast growth in isolation, each allele was cloned into the pAG425 vector and transformed into the ptetO7-*ALA1* yeast strain, followed by a complementation growth assay as described above. Interestingly, none of the deletions fully complemented loss of *ALA1* (Figure 3B), indicating that the affected residues are indeed important for AARS1 protein function. To determine if the engineered mutations impact protein stability, each allele was evaluated for an effect on AARS1 protein levels via western blot analysis. The N944_G946del allele led to no detectable AARS1 protein (Figure 3C), providing an explanation for its failure to complement in yeast. The C947 deletion significantly reduced AARS1 expression (Figure 3C), but still showed partial complementation in yeast (Figure 3B), suggesting that this residue may be more important for stability than for AARS1 function. The deletions K943_Q949del and Q949_E950del similarly reduced AARS1 levels (Figure 3C) and led to less yeast growth than C947del (Figure 3B), suggesting that these deletions reduce function as well as protein abundance. Only the globular domain deletion Q855* resulted in a complete loss of yeast cell growth (Figure 3B) with no detectable decrease in protein levels (Figure 3C). Additionally, this mutation is not predicted to significantly alter the structure of the protein outside of the dimerization domain (Supplemental Figure 5). Based on these data we concluded that Q855* is the most appropriate allele to test for reduced dimerization.

To test if Q855* reduces binding to wild-type AARS1, HEK293T cells were transfected with wild-type AARS1-3xFLAG and either wild-type or Q855* AARS1-6xHis. First, all transfections resulted in robust AARS1 protein expression (Figure 4A), consistent with Q855* AARS1 not significantly impacting protein levels. Second, immunoprecipitation of wild-type AARS1-6xHis co-precipitated wild-type AARS1-3xFLAG, confirming an interaction between the two tagged wild-type subunits (Figure 4B). However, immunoprecipitation for Q855* AARS1-6xHis did not co-precipitate wild-type AARS1-3xFLAG (Figure 4B), indicating that Q855* reduces or ablates binding to the wild-type AARS1 protein. These findings were supported by performing the reciprocal experiment. Here, immunoprecipitation of wild-type AARS1-3xFLAG co-immunoprecipitated wild-type AARS1-6xHis, but did not co-immunoprecipitate Q855* AARS1-6xHis (Figure 4C). These data demonstrate that the engineered Q855* *AARS1* variant reduces the interaction with wild-type AARS1, and for the first time experimentally confirms that the C-terminal globular domain is required for dimerization.

**Figure 4.**
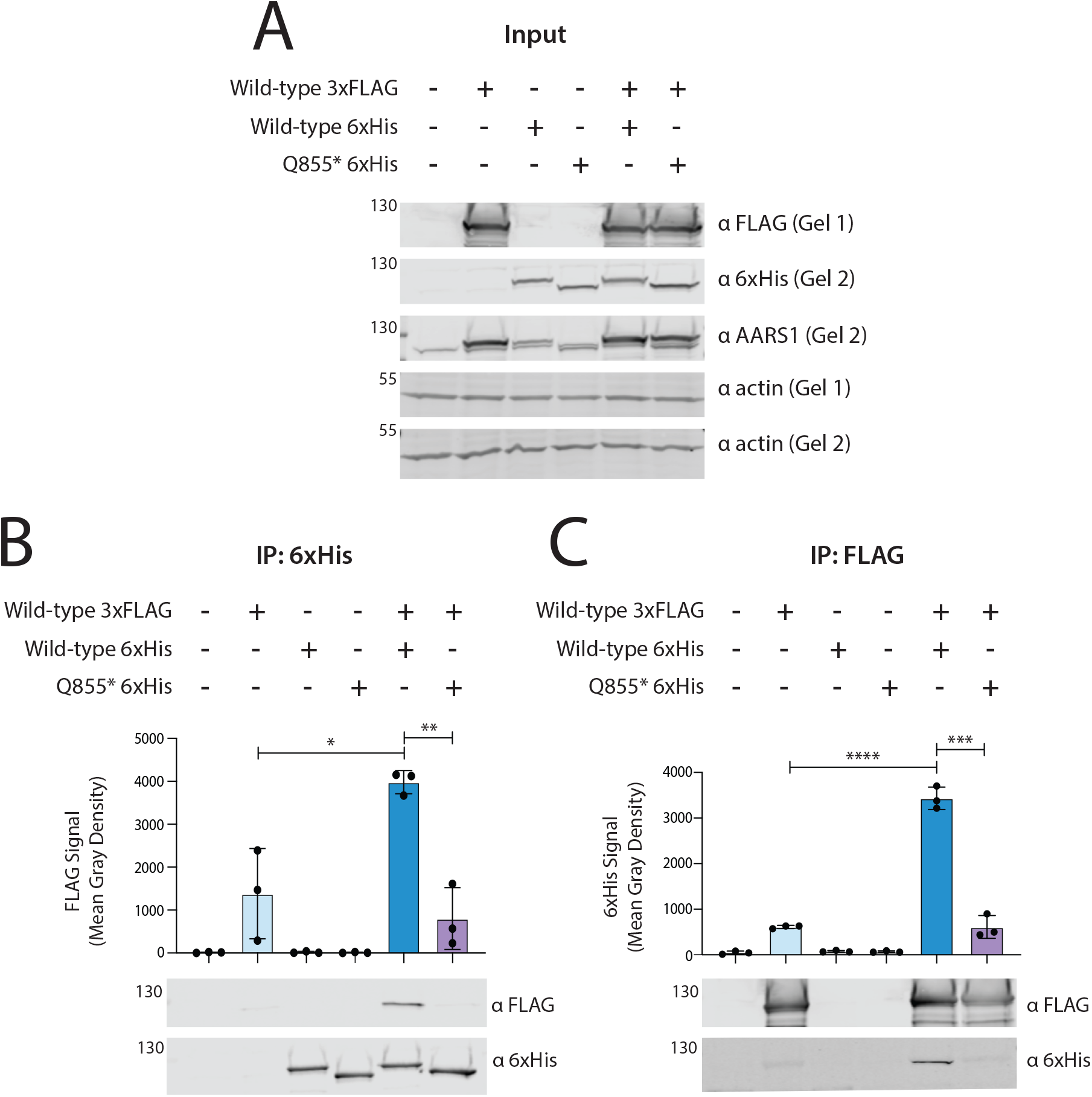
Q855* AARS1 impairs dimerization with wild-type AARS1. **(A)** HEK293T cells were transfected with vectors to co-express wild-type or Q855* human *AARS1*, and a western blot was performed to detect the resulting proteins, as well as endogenous loading controls. The image is representative of three independent replicates. The presence or absence of each construct is shown across the top of the image, protein molecular weights are indicated in kilodaltons (kDa) along the left side, and antibodies are indicated along the right side. **(B)** After immunoprecipitation with an anti-6xHis antibody, a western blot was performed to detect co-immunoprecipitated proteins. The presence or absence of each construct is indicated at the top of the panel, and a representative image from three independent replicates is shown at the bottom of the panel (with annotation as in panel A). The middle of the panel shows ImageJ quantification of 3xFLAG-tagged AARS1 signal. **(C)** After immunoprecipitation with an anti-FLAG antibody, a western blot was performed to detect co-immunoprecipitated proteins. The panel is organized similar to panel B, with ImageJ quantification of 6xHis-tagged AARS1 signal in the middle of the panel. For the ImageJ quantification shown in panel B and panel C, bars indicate the mean value and one standard deviation of three replicates. Unpaired t-tests were performed to determine if the difference in signal intensity between samples was statistically significant. **** p<0.0001, *** p<0.001, ** p<0.01, * p<0.05.

### Reducing the dimerization capacity of pathogenic *AARS1* alleles rescues yeast growth

If G102R and R329H *AARS1* act as dominant-negative alleles to impair yeast growth, then reducing the dimerization between mutant and wild-type AARS1 proteins should rescue yeast growth. To test this, the Q855* mutation was introduced in *cis* with either G102R or R329H using site-directed mutagenesis. These double mutants were then cloned into pAG425 and transformed into the ptetO7-*ALA1* strain. Both G102R+Q855* *AARS1* and R329H+Q855* *AARS1* produced a truncated AARS1 protein at levels comparable to wild-type AARS1 (Supplemental Figure 6A and B).

Complementation assays expressing G102R+Q855* *AARS1* or R329H+Q855* *AARS1* in the presence of an empty p413 vector showed no yeast growth, consistent with the double-mutants acting as loss-of-function alleles (Figure 5A). These alleles were then tested in the presence of wild-type AARS1 expressed from the p413 vector. As before, neither the control allele G757* *AARS1* nor wildtype *AARS1* repressed yeast growth, and both G102R and R329H repressed yeast growth (Figure 5B, 5C). Importantly, placing Q855* in *cis* with either G102R or R329H ameliorated the growth phenotype and increased the mean growth 24-fold or 82-fold, respectively. (Figure 5B, 5C). This rescued growth was comparable to that of yeast expressing the G757* control allele or wild-type *AARS1* (Figure 5C). These data demonstrate that disrupting the dimerization of G102R or R329H with wild-type AARS1 is sufficient to rescue the dominant growth phenotype, and show that the reduced growth phenotype is a result of mutant AARS1 dimerizing with wild-type AARS1. In sum, these data provide evidence that neuropathy-associated *AARS1* alleles are loss-of-function variants that dominantly repress yeast growth through dimerization with the wild-type subunit; *i.e*. that they act via a dominant-negative mechanism.

**Figure 5.**
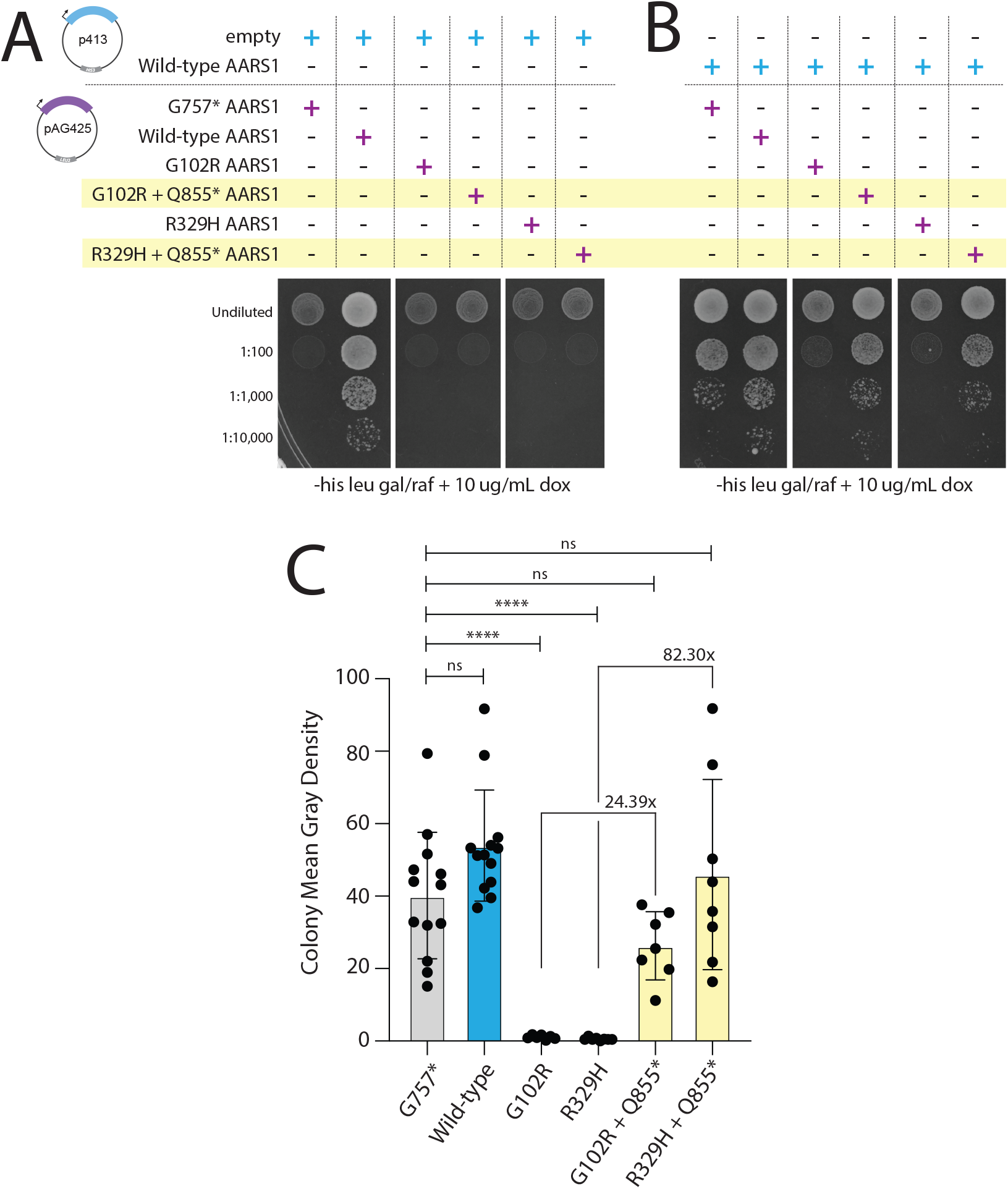
Reducing dimerization of G102R and R329H with wild-type AARS1 rescues yeast growth. (**A**) Yeast with the doxycycline-repressible endogenous *ALA1* locus were transformed with an empty p413 vector and a pAG425 vector expressing wild-type or mutant human *AARS1*. Cultures were plated undiluted or diluted on media lacking histidine and leucine, and containing galactose/raffinose and doxycycline. (**B**) Similar to strains shown in panel A, except that yeast were transformed with p413 expressing wild-type human AARS1. For both panels, the vectors present in each experiment are indicated across the top, the dilution of the spotted yeast cultured is indicated on the left, and the media conditions are indicated across the bottom (his = histidine; leu = leucine; gal = galactose; raf = raffinose; dox = doxycycline). **(C)** Yeast spot intensity was quantified using ImageJ. Bars represent the mean and one standard deviation. Thirteen biological replicates were assessed for G757* and wild-type *AARS1*, eight for R329H and R329H+Q855*, and seven for G102R and G102R+Q855*. The indicated fold-change between the G102R strain and the G102R+Q855* strain, and the fold-change between the R329H strain and the R329H+Q855* strain, were both calculated using the mean of each sample. To compare yeast growth to the strain expressing both wild-type and G757* *AARS1*, a oneway ANOVA with Dunnett’s multiple comparisons test was performed. **** p<0.0001, *** p<0.001, ns=not significant.

### Sequences that encode the AARS1 anticodon-binding domain are susceptible to recurrent dominant-negative mutations

If G102R and R329H *AARS1* are dominant-negative alleles, it is likely that other *AARS1* variants found in patients with dominant CMT are also dominant-negative alleles. To investigate this possibility, we focused on the R329 residue and the surrounding *AARS1* coding region. Previous work by McLaughlin et al. identified a high degree of cytosine methylation in this area, making it susceptible to cytosine deamination. This study predicted numerous missense variants that could arise from such CàT changes (42). One such predicted change, R326W, was later found in a multi-generational family with CMT (23). Here, we report four additional individuals with mutations at the R329 residue, all of whom were examined by a neurologist and presented with axonal neuropathy (Supplemental Table 1). One individual is heterozygous for R329S, a variant previously predicted by the cytosine methylation analysis (42). The other three individuals comprise the eleventh family with CMT caused by the R329H variant, further strengthening the argument that this is a pathogenic allele and that the R329 residue is subject to recurrent mutation. In addition to R329H and R329S, McLaughlin et al. predicted that R329C could arise at this residue; however, this allele has yet to be identified in patients with CMT disease. Therefore, we assessed all three variants (R326W, R329S, and R329C) in our humanized yeast dominant-negative assay.

The R326W, R329S, and R329C variants were introduced into the *AARS1* open reading frame with site-directional mutagenesis, then cloned into the galactose-inducible pAG425 expression vector and transformed into the ptetO7-*ALA1* yeast strain. G757* and wild-type *AARS1* were included as controls. To determine if our yeast model can distinguish between pathogenic and non-pathogenic *AARS1* alleles, we also tested G931S *AARS1*, which is a benign polymorphism found in the general population (with a gnomAD allele count of 2,147 out of 282,842 chromosomes including 20 homozygous individuals [52]). As previously reported (23), R326W did not support yeast growth in the absence of *ALA1* (Figure 6A). Consistent with the functional importance of this region, R329S and R329C also did not support yeast growth (Figure 6A). These three variants were then tested in the presence of the wild-type *AARS1* allele expressed from p413. R326W, R329S, and R329C *AARS1* all repressed yeast growth in the presence of wild-type *AARS1*, compared to G757*, wildtype, or G931S *AARS1* (Figure 6B). Notably, as was the case for G102R and R329H, this growth defect was improved by restoring the expression of endogenous *ALA1* (Supplemental Figure 3C, 3D), supporting the argument that the phenotype is due to an alanine-tRNA charging defect.

**Figure 6.**
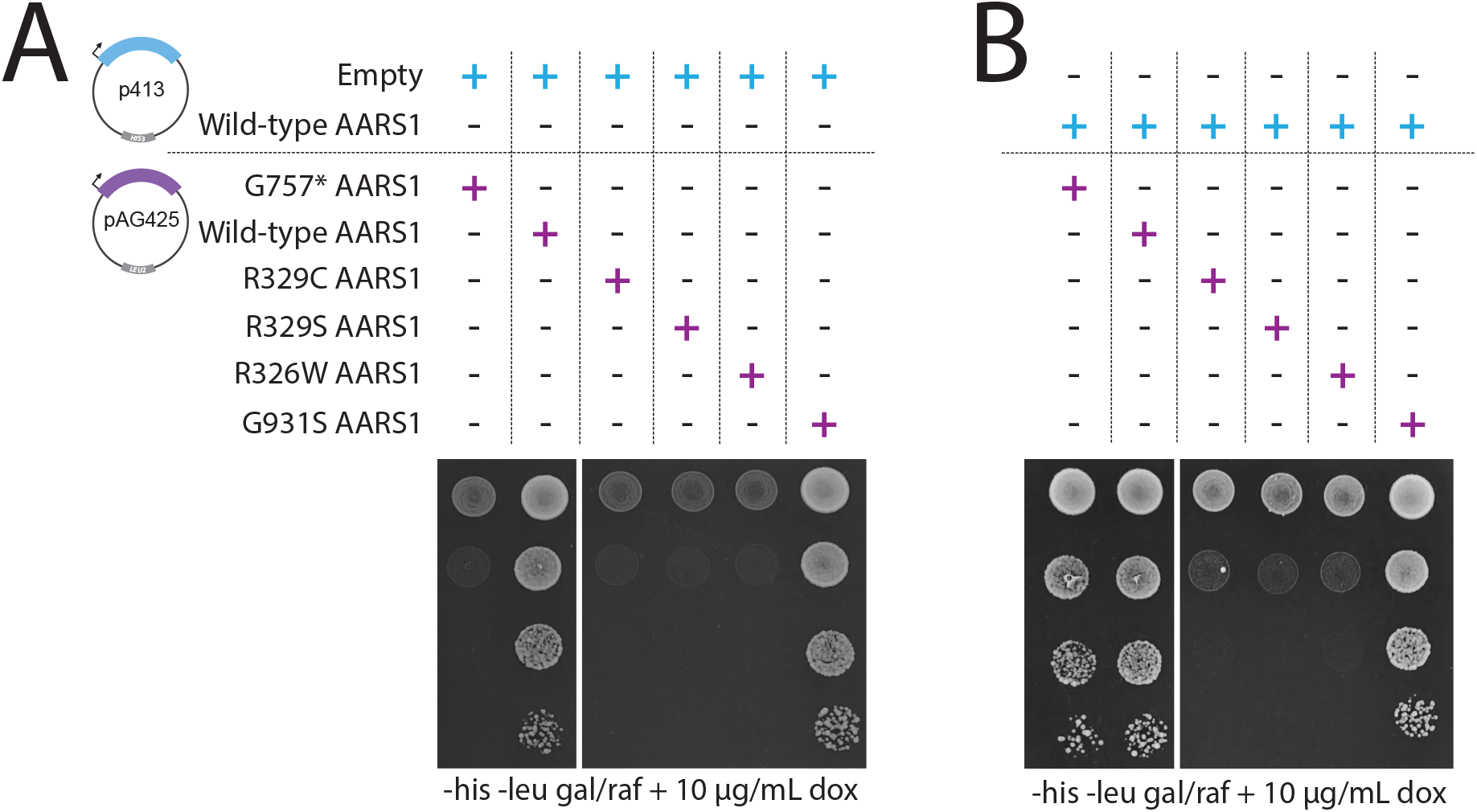
R329C, R329S, and R326W *AARS1* are loss-of-function alleles that dominantly repress yeast cell growth. **(A)** Yeast with the doxycycline-repressible endogenous *ALA1* locus were transformed with a p413 vector with no insert, and a pAG425 vector expressing wild-type or mutant *AARS1*. Yeast cultures were spotted undiluted or diluted on media lacking histidine and leucine, and containing galactose/raffinose and doxycycline. **(B)** Similar experiment to that described in panel A, except that yeast express wild-type human *AARS1* from p413. For both panels, the vectors present in each experiment are indicated across the top, the dilution of the spotted yeast cultured is indicated on the left, and the media conditions are shown across the bottom (his = histidine; leu = leucine; gal = galactose; raf = raffinose; dox = doxycycline). Images are representative of three replicates.

To determine if the dominant growth repression associated with these three alleles depends on their ability to dimerize with wild-type AARS1, the dimer-disrupting Q855* variant was introduced *in cis* with either R326W, R329S, or R329C. These double-mutant alleles were transformed into yeast expressing an empty p413 vector, or expressing wild-type *AARS1* from p413. Strains were then plated on galactose (to express the double-mutant *AARS1* allele) and doxycycline (to repress endogenous *ALA1*). In the presence of the empty p413 vector, the double-mutants showed no yeast growth, consistent with the double-mutants acting as loss-of-function alleles (Figure 7A). However, when co-expressed with wild-type *AARS1*, Q855* *in cis* with R329C, R329S, or R326W rescued the repressed yeast growth for all three variants, increasing growth by 15-fold, 26-fold, and 29-fold respectively (Figure 7B, 7C). The growth of the double-mutants was not significantly different than the growth of the yeast expressing the control allele G757* or wild-type *AARS1* (Figure 7C). Combined, these data indicate that R326W, R329S, and R329C are dominant-negative alleles. Together with G102R and R329H, this study presents evidence that neuropathy-associated *AARS1* variants act as dominant-negative alleles.

**Figure 7.**
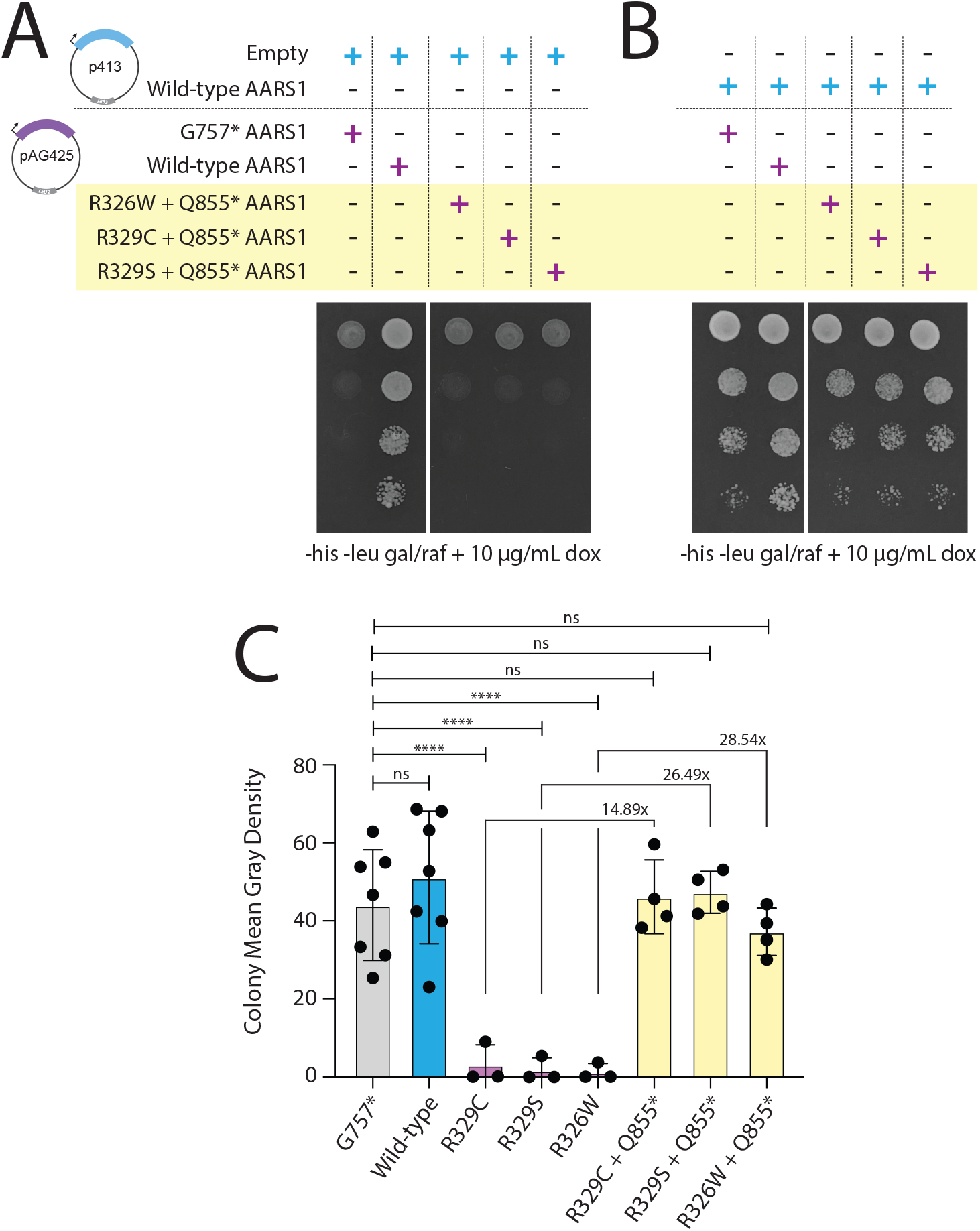
Reducing dimerization of R329C, R329S, or R326W with wild-type AARS1 rescues yeast growth. **(A)** Yeast with the doxycycline-repressible *ALA1* locus were transformed with an empty p413 vector and a pAG425 vector expressing either wild-type or mutant *AARS1*. Yeast were spotted undiluted or diluted on media lacking histidine and leucine, and containing galactose/raffinose and doxycycline. **(B)** An experiment similar to that described in panel A, except that yeast express wildtype human *AARS1* from the p413 vector. **(C)** Yeast spot intensity was quantified using ImageJ analysis; bars represent the mean and one standard deviation. At least three biological replicates were assessed for all variants. The indicated fold change between the strains expressing R329C, R329S, or R326W, and their counterpart with Q855* *in cis*, was calculated using the mean intensity of each condition. To compare yeast growth to that of the strain expressing both wild-type and G757* *AARS1*, a one-way ANOVA with Dunnett’s multiple comparisons test was performed. ****p<0.0001, ns=not significant.

## DISCUSSION

In this study, we developed a humanized yeast assay to study the molecular mechanism of neuropathy-associated *AARS1* alleles. We demonstrate that multiple pathogenic, loss-of-function *AARS1* variants repress yeast growth when co-expressed with the wild-type *AARS1* allele. We also show that these variants retain the ability to dimerize with the wild-type AARS1 protein, and that disrupting this interaction by deleting critical dimerization residues from the mutant protein is sufficient to rescue the repressed yeast growth. These data provide evidence that neuropathy-associated *AARS1* alleles act via a dominant-negative mechanism that represses the activity of wild-type AARS1.

It will be important to employ the dominant-negative yeast model reported here to study other variants in *AARS1*. These studies will enable rapid characterization of newly identified *AARS1* alleles and will provide functional evidence supporting pathogenicity. Furthermore, an unbiased, massively parallel mutagenesis approach utilizing this model will help predict dominant-negative *AARS1* alleles that may be identified in patients in the future, such as R329C. This yeast model can also be adapted to study and predict pathogenic variants in *GARS1, HARS1, YARS1*, and *WARS1*. We hypothesize that dominant pathogenic variants in these other neuropathy-associated ARS genes will show dominant-negative effects, because these genes also encode homo-dimeric ARS enzymes (53–56). Classifying pathogenic variants in these additional ARS loci as dominant-negative alleles will also contribute to defining a common mechanism of ARS-mediated dominant neuropathy, and will provide critical insight into how missense ARS variants lead to depleted pools of charged tRNA, which ultimately impairs protein translation (36, 38–41).

While the data presented here strongly argue in favor of a dominant-negative effect, it is possible that a dominant-negative and neomorphic gain-of-function effects work in concert to exacerbate neuronal pathology. A recent study showed that the pathogenic *AARS1* variants N71Y, G102R, and R329H each cause a conformational change that enables binding to Neuropilin-1 (32), a widely expressed receptor that modulates a variety of signalling pathways and that is critical for neurovascular development (57). Although such an interaction might compound the damage in patient neurons, our yeast model demonstrates that a neuron-specific (or mammalian-specific) interaction is not required for pathogenic *AARS1* alleles to exert dominant effects.

One limitation of our yeast model is that the allelic expression is intentionally skewed, with the pathogenic allele over-expressed relative to the wild-type allele. This does not accurately reflect the presumably equal expression of wild-type and mutant alleles in the tissues of a heterozygous patient. Therefore, any dominant-negative effects in patients are likely to be weaker than those demonstrated here, and less likely to have such dramatic consequences for cell viability. Indeed, such a tempered effect in patient cells is more consistent with the late-onset and tissue-restricted patient phenotype. A terminally differentiated peripheral neuron that must maintain local protein translation far from the soma may be particularly susceptible to even mild dominant-negative effects of an ARS mutation. To determine if a dominant-negative effect drives dominant *AARS1*-mediated peripheral neuropathy, a knock-in animal model (*e.g*., mouse or worm) with an axonal pathology is required. Then, a dimer-disrupting variant such as Q855* can be introduced *in cis* with the pathogenic allele to determine if this ameliorates the neuronal phenotype.

Although there has been debate (32, 51) as to whether AARS1 functions as a dimer or a monomer, the results of our study demonstrate that it likely functions as a dimer *in vivo*, which is a requirement for a dominant-negative mechanism of disease. Previous studies using purified AARS1 protein for *in vitro* enzymatic assays determined that the C-terminal dimerization domain (beginning at G757) is dispensable for aminoacylation (51). However, our data demonstrate that, at least in a cellular context, this domain is likely required for protein stability; truncating the protein at this residue leads to undetectable levels of AARS1 protein when this allele is expressed in yeast cells. Furthermore, the globular domain beginning at Q855*, while not required to maintain protein levels in yeast or mammalian cells, is required for both AARS1 dimerization and activity in yeast. This indicates that dimerization is necessary for AARS1 function *in vivo*. Finally, chemically crosslinked AARS1 from human fibroblast cells is detected at two molecular weights, one corresponding to a monomeric form and one corresponding to a dimeric form. Overall, these data reveal that AARS1 dimerization is important for protein function *in vivo*.

To the best of our knowledge, there have been few yeast systems developed to test human pathogenic variants for a dominant-negative effect. Notable examples include reporter assays to test for dominant-negative p53 mutations (58, 59) and enzymatic evaluation of yeast expressing dominant-negative UDP-galactose-4-epimerase (GALE) alleles (60, 61). In general, humanized yeast approaches are limited by the ability to recapitulate mammalian gene function in yeast cells. As such, they are best suited to study variants in genes with highly conserved functions, particularly ones that impact a readily observable phenotype, such as cellular growth. Using strategies similar to those presented here, variants in genes that fit these requirements can be efficiently interrogated for loss-of-function and/or dominant-negative properties, providing critical data to aid in patient diagnosis and further defining how genetic variation impacts gene function.

Here, we describe a tractable yeast model for rapidly evaluating patient variants in aminoacyl-tRNA synthetase genes for a dominant-negative effect. We demonstrate that multiple well-characterized, pathogenic *AARS1* variants repress yeast growth when co-expressed with wild-type *AARS1*, but that this phenotype is rescued when the mutant subunit is prevented from dimerizing with the wild-type subunit. This work presents the first direct evidence that pathogenic ARS alleles can exert a dominant-negative effect. This mechanism may at least partly contribute to the effect of dominant pathogenic ARS variants in reducing the availability of charged tRNA, causing ribosome stalling, and triggering the integrated stress response (as recently shown for several *GARS1* alleles [38, 40, 41]). In sum, this work contributes to further delineating a loss-of-function mechanism of ARS-mediated dominant peripheral neuropathy.

## DATA AVAILABILITY

Not applicable.

## SUPPLEMENTARY DATA

Supplementary Data are available online.

## ACKNOWLEDGEMENT

R.M. is supported by the Michigan Pre-doctoral Training in Genetics Program (GM007544) and an individual National Research Service Award (NRSA) from the National Institute of Neurological Diseases and Stroke (NS108510). A.A. is supported by a grant from the National Institute of General Medical Sciences (GM136441). K.S.K. and T.J.S. are supported by a grant from the National Institute of General Medical Sciences (GM128836). The authors would like to thank: Dr. Thomas Glover and Dr. Miriam Meisler for their gifts of fibroblast cell lines from unaffected individuals; Dr. Thomas Wilson for advice on yeast genetic experimentation; Dr. John Moldovan, Dr. Trenton Frisbie, and Dr. Stephanie Moon for advice on protein-protein interaction experiments; and Dr. John Moran for advice on protein biochemistry and yeast genetic studies, and for frequent discussions on the goals of the study. The authors would also like to thank Alex Mark for his help running AlphaFold Colab, and Dr. Laura Lavery for her advice on predicting protein structure.

## REFERENCES

1. Shy, M.E., Lupski, J.R., Chance, P.F., Klein, C.J. and Dyck, P.J. Hereditary Motor and Sensory Neuropathies: An Overview of Clinical, Genetic, Electrophysiologic, and Pathologic Features.

2. Scherer, S.S., Kleopa, K.A. and Benson, M.D. (2020) Chapter 21 - Peripheral neuropathies. In Rosenberg, R.N., Pascual, J.M. (eds), Rosenberg’s Molecular and Genetic Basis of Neurological and Psychiatric Disease (Sixth Edition). Academic Press, pp. 345–375.

3. Skre, H. (1974) Genetic and clinical aspects of Charcot-Marie-Tooth’s disease. Clinical Genetics, 6, 98–118.

4. Braathen, G.J., Sand, J.C., Lobato, A., Høyer, H. and Russell, M.B. (2011) Genetic epidemiology of Charcot-Marie-Tooth in the general population. Eur J Neurol, 18, 39–48.

5. Laura, M., Pipis, M., Rossor, A.M. and Reilly, M.M. (2019) Charcot-Marie-Tooth disease and related disorders: an evolving landscape. [Miscellaneous Article]. Current Opinion in Neurology, 32, 641–650.

6. Antonellis, A. and Green, E.D. (2008) The role of aminoacyl-tRNA synthetases in genetic diseases. Annual review of genomics and human genetics, 9, 87–107.

7. Latour, P., Thauvin-Robinet, C., Baudelet-Méry, C., Soichot, P., Cusin, V., Faivre, L., Locatelli, M.-C., Mayençon, M., Sarcey, A., Broussolle, E., et al. (2010) A Major Determinant for Binding and Aminoacylation of tRNAAla in Cytoplasmic Alanyl-tRNA Synthetase Is Mutated in Dominant Axonal Charcot-Marie-Tooth Disease. The American Journal of Human Genetics, 86, 77–82.

8. Safka Brozkova, D., Deconinck, T., Griffin, L.B., Ferbert, A., Haberlova, J., Mazanec, R., Lassuthova, P., Roth, C., Pilunthanakul, T., Rautenstrauss, B., et al. (2015) Loss of function mutations in HARS cause a spectrum of inherited peripheral neuropathies. Brain : a journal of neurology, 138, 2161–2172.

9. Antonellis, A., Ellsworth, R.E., Sambuughin, N., Puls, I., Abel, A., Lee-Lin, S.-Q., Jordanova, A., Kremensky, I., Christodoulou, K., Middleton, L.T., et al. (2003) Glycyl tRNA synthetase mutations in Charcot-Marie-Tooth disease type 2D and distal spinal muscular atrophy type V. The American Journal of Human Genetics, 72, 1293–1299.

10. Tsai, P.-C., Soong, B.-W., Mademan, I., Huang, Y.-H., Liu, C.-R., Hsiao, C.-T., Wu, H.-T., Liu, T.-T., Liu, Y.-T., Tseng, Y.-T., et al. (2017) A recurrent WARS mutation is a novel cause of autosomal dominant distal hereditary motor neuropathy. Brain : a journal of neurology.

11. Jordanova, A., Irobi, J., Thomas, F.P., Van Dijck, P., Meerschaert, K., Dewil, M., Dierick, I., Jacobs, A., De Vriendt, E., Guergueltcheva, V., et al. (2006) Disrupted function and axonal distribution of mutant tyrosyl-tRNA synthetase in dominant intermediate Charcot-Marie-Tooth neuropathy. Nature Genetics, 38, 197–202.

12. Gonzalez, M., McLaughlin, H., Houlden, H., Guo, M., Yo-Tsen, L., Hadjivassilious, M., Speziani, F., Yang, X.-L., Antonellis, A., Reilly, M.M., et al. (2013) Exome sequencing identifies a significant variant in methionyl-tRNA synthetase (*MARS*) in a family with late-onset CMT2. J Neurol Neurosurg Psychiatry, 84, 1247–1249.

13. Gillespie, M.K., McMillan, H.J., Kernohan, K.D., Pena, I.A., Meyer-Schuman, R., Antonellis, A. and Boycott, K.M. A Novel Mutation in MARS in a Patient with Charcot-Marie-Tooth Disease, Axonal, Type 2U with Congenital Onset. J Neuromuscul Dis, 6, 333–339.

14. Meyer-Schuman, R. and Antonellis, A. (2017) Emerging mechanisms of aminoacyl-tRNA synthetase mutations in recessive and dominant human disease. Human molecular genetics, 26, R114–R127.

15. Kuo, M.E. and Antonellis, A. (2020) Ubiquitously Expressed Proteins and Restricted Phenotypes: Exploring Cell-Specific Sensitivities to Impaired tRNA Charging. Trends in Genetics, 36, 105–117.

16. Kuo, M.E., Theil, A.F., Kievit, A., Malicdan, M.C., Introne, W.J., Christian, T., Verheijen, F.W., Smith, D.E.C., Mendes, M.I., Hussaarts-Odijk, L., et al. (2019) Cysteinyl-tRNA Synthetase Mutations Cause a Multi-System, Recessive Disease That Includes Microcephaly, Developmental Delay, and Brittle Hair and Nails. Am. J. Hum. Genet., 104, 520–529.

17. McLaughlin, H.M., Sakaguchi, R., Liu, C., Igarashi, T., Pehlivan, D., Chu, K., Iyer, R., Cruz, P., Cherukuri, P.F., Hansen, N.F., et al. (2010) Compound heterozygosity for loss-of-function lysyl-tRNA synthetase mutations in a patient with peripheral neuropathy. Am J Hum Genet, 87, 560–566.

18. Manole, A., Efthymiou, S., O’Connor, E., Mendes, M.I., Jennings, M., Maroofian, R., Davagnanam, I., Mankad, K., Lopez, M.R., Salpietro, V., et al. (2020) De Novo and Bi-allelic Pathogenic Variants in NARS1 Cause Neurodevelopmental Delay Due to Toxic Gain-of-Function and Partial Loss-of-Function Effects. Am J Hum Genet, 107, 311–324.

19. Seburn, K.L., Nangle, L.A., Cox, G.A., Schimmel, P. and Burgess, R.W. (2006) An active dominant mutation of glycyl-tRNA synthetase causes neuropathy in a Charcot-Marie-Tooth 2D mouse model. Neuron, 51, 715–726.

20. Achilli, F., Bros-Facer, V., Williams, H.P., Banks, G.T., AlQatari, M., Chia, R., Tucci, V., Groves, M., Nickols, C.D., Seburn, K.L., et al. (2009) An ENU-induced mutation in mouse glycyl-tRNA synthetase (GARS) causes peripheral sensory and motor phenotypes creating a model of Charcot-Marie-Tooth type 2D peripheral neuropathy. Disease models & mechanisms, 2, 359–373.

21. Morelli, K.H., Griffin, L.B., Pyne, N.K., Wallace, L.M., Fowler, A.M., Oprescu, S.N., Takase, R., Wei, N., Meyer-Schuman, R., Mellacheruvu, D., et al. (2019) Allele-specific RNA interference prevents neuropathy in Charcot-Marie-Tooth disease type 2D mouse models. J. Clin. Invest., 129, 5568–5583.

22. Oprescu, S.N., Griffin, L.B., Beg, A.A. and Antonellis, A. (2017) Predicting the pathogenicity of aminoacyl-tRNA synthetase mutations. Methods, 113, 139–151.

23. Weterman, M.A.J., Kuo, M., Kenter, S.B., Gordillo, S., Karjosukarso, D.W., Takase, R., Bronk, M., Oprescu, S., van Ruissen, F., Witteveen, R.J.W., et al. (2018) Hypermorphic and hypomorphic AARS alleles in patients with CMT2N expand clinical and molecular heterogeneities. Hum. Mol. Genet., 27, 4036–4050.

24. Bansagi, B., Antoniadi, T., Burton-Jones, S., Murphy, S.M., McHugh, J., Alexander, M., Wells, R., Davies, J., Hilton-Jones, D., Lochmüller, H., et al. (2015) Genotype/phenotype correlations in AARS-related neuropathy in a cohort of patients from the United Kingdom and Ireland. J Neurol, 262, 1899–1908.

25. Lee, D.C., Meyer-Schuman, R., Bacon, C., Shy, M.E., Antonellis, A. and Scherer, S.S. (2019) A recurrent GARS mutation causes distal hereditary motor neuropathy. J Peripher Nerv Syst, 24, 320–323.

26. Markovitz, R., Ghosh, R., Kuo, M.E., Hong, W., Lim, J., Bernes, S., Manberg, S., Crosby, K., Tanpaiboon, P., Bharucha-Goebel, D., et al. (2020) GARS-related disease in infantile spinal muscular atrophy: Implications for diagnosis and treatment. American Journal of Medical Genetics Part A, 182, 1167–1176.

27. Abbott, J.A., Meyer-Schuman, R., Lupo, V., Feely, S., Mademan, I., Oprescu, S.N., Griffin, L.B., Alberti, M.A., Casasnovas, C., Aharoni, S., et al. (2018) Substrate interaction defects in histidyl-tRNA synthetase linked to dominant axonal peripheral neuropathy. Hum. Mutat., 39, 415–432.

28. Wang, B., Li, X., Huang, S., Zhao, H., Liu, J., Hu, Z., Lin, Z., Liu, L., Xie, Y., Jin, Q., et al. (2019) A novel WARS mutation (p.Asp314Gly) identified in a Chinese distal hereditary motor neuropathy family. Clinical Genetics, 96, 176–182.

29. He, W., Bai, G., Zhou, H., Wei, N., White, N.M., Lauer, J., Liu, H., Shi, Y., Dumitru, C.D., Lettieri, K., et al. (2015) CMT2D neuropathy is linked to the neomorphic binding activity of glycyl-tRNA synthetase. Nature, 526, 710–714.

30. Sleigh, J.N., Dawes, J.M., West, S.J., Wei, N., Spaulding, E.L., Gómez-Martín, A., Zhang, Q., Burgess, R.W., Cader, M.Z., Talbot, K., et al. (2017) Trk receptor signaling and sensory neuron fate are perturbed in human neuropathy caused by Gars mutations. Proceedings of the National Academy of Sciences of the United States of America.

31. Bervoets, S., Wei, N., Erfurth, M.-L., Yusein-Myashkova, S., Ermanoska, B., Mateiu, L., Asselbergh, B., Blocquel, D., Kakad, P., Penserga, T., et al. (2019) Transcriptional dysregulation by a nucleus-localized aminoacyl-tRNA synthetase associated with Charcot-Marie-Tooth neuropathy. Nat Commun, 10, 5045.

32. Sun, L., Wei, N., Kuhle, B., Blocquel, D., Novick, S., Matuszek, Z., Zhou, H., He, W., Zhang, J., Weber, T., et al. (2021) CMT2N-causing aminoacylation domain mutants enable Nrp1 interaction with AlaRS. Proc Natl Acad Sci U S A, 118.

33. Mo, Z., Zhao, X., Liu, H., Hu, Q., Chen, X.-Q., Pham, J., Wei, N., Liu, Z., Zhou, J., Burgess, R.W., et al. (2018) Aberrant GlyRS-HDAC6 interaction linked to axonal transport deficits in Charcot-Marie-Tooth neuropathy. Nat Commun, 9, 1007.

34. Antonellis, A., Lee-Lin, S.-Q., Wasterlain, A., Leo, P., Quezado, M., Goldfarb, L.G., Myung, K., Burgess, S., Fischbeck, K.H. and Green, E.D. (2006) Functional analyses of glycyl-tRNA synthetase mutations suggest a key role for tRNA-charging enzymes in peripheral axons. The Journal of neuroscience : the official journal of the Society for Neuroscience, 26, 10397–10406.

35. Motley, W.W., Seburn, K.L., Nawaz, M.H., Miers, K.E., Cheng, J., Antonellis, A., Green, E.D., Talbot, K., Yang, X.-L., Fischbeck, K.H., et al. (2011) Charcot-Marie-Tooth–Linked Mutant GARS Is Toxic to Peripheral Neurons Independent of Wild-Type GARS Levels. PLoS Genet, 7.

36. Mullen, P., Abbott, J.A., Wellman, T., Aktar, M., Fjeld, C., Demeler, B., Ebert, A.M. and Francklyn, C.S. (2020) Neuropathy-associated histidyl-tRNA synthetase variants attenuate protein synthesis in vitro and disrupt axon outgrowth in developing zebrafish. FEBS J., 10.1111/febs.15449.

37. Nangle, L.A., Zhang, W., Xie, W., Yang, X.-L. and Schimmel, P. (2007) Charcot-Marie-Tooth disease-associated mutant tRNA synthetases linked to altered dimer interface and neurite distribution defect. Proceedings of the National Academy of Sciences, 104, 11239–11244.

38. Mendonsa, S., von Kuegelgen, N., Bujanic, L. and Chekulaeva, M. (2021) Charcot–Marie–Tooth mutation in glycyl-tRNA synthetase stalls ribosomes in a pre-accommodation state and activates integrated stress response. Nucleic Acids Research, 10.1093/nar/gkab730.

39. Niehues, S., Bussmann, J., Steffes, G., Erdmann, I., Köhrer, C., Sun, L., Wagner, M., Schäfer, K., Wang, G., Koerdt, S.N., et al. (2015) Impaired protein translation in Drosophila models for Charcot–Marie–Tooth neuropathy caused by mutant tRNA synthetases. Nature Communications, 6, 7520–13.

40. Spaulding, E.L., Hines, T.J., Bais, P., Tadenev, A.L.D., Schneider, R., Jewett, D., Pattavina, B., Pratt, S.L., Morelli, K.H., Stum, M.G., et al. (2021) The integrated stress response contributes to tRNA synthetase–associated peripheral neuropathy. Science, 373, 1156–1161.

41. Zuko, A., Mallik, M., Thompson, R., Spaulding, E.L., Wienand, A.R., Been, M., Tadenev, A.L.D., van Bakel, N., Sijlmans, C., Santos, L.A., et al. (2021) tRNA overexpression rescues peripheral neuropathy caused by mutations in tRNA synthetase. Science, 373, 1161–1166.

42. McLaughlin, H.M., Sakaguchi, R., Giblin, W., Wilson, T.E., Biesecker, L., Lupski, J.R., Talbot, K., Vance, J.M., Züchner, S., Lee, Y.-C., et al. (2012) A Recurrent loss-of-function alanyl-tRNA synthetase (AARS) mutation in patients with charcot-marie-tooth disease type 2N (CMT2N). Human mutation, 33, 244–253.

43. Lee, A.J., Nam, D.E., Choi, Y.J., Nam, S.H., Choi, B.-O. and Chung, K.W. (2020) Alanyl-tRNA synthetase 1 (AARS1) gene mutation in a family with intermediate Charcot-Marie-Tooth neuropathy. Genes Genomics, 42, 663–672.

44. Motley, W.W., Griffin, L.B., Mademan, I., Baets, J., De Vriendt, E., De Jonghe, P., Antonellis, A., Jordanova, A. and Scherer, S.S. (2015) A novel AARS mutation in a family with dominant myeloneuropathy. Neurology, 84, 2040–2047.

45. Petropavlovskiy, A.A., Tauro, M.G., Lajoie, P. and Duennwald, M.L. (2020) A Quantitative Imaging-Based Protocol for Yeast Growth and Survival on Agar Plates. STAR Protoc, 1, 100182.

46. Jumper, J., Evans, R., Pritzel, A., Green, T., Figurnov, M., Ronneberger, O., Tunyasuvunakool, K., Bates, R., Žídek, A., Potapenko, A., et al. (2021) Highly accurate protein structure prediction with AlphaFold. Nature, 596, 583–589.

47. Mnaimneh, S., Davierwala, A.P., Haynes, J., Moffat, J., Peng, W.-T., Zhang, W., Yang, X., Pootoolal, J., Chua, G., Lopez, A., et al. (2004) Exploration of Essential Gene Functions via Titratable Promoter Alleles. Cell, 118, 31–44.

48. Mumberg, D., Müller, R. and Funk, M. (1995) Yeast vectors for the controlled expression of heterologous proteins in different genetic backgrounds. Gene, 156, 119–122.

49. Alberti, S., Gitler, A.D. and Lindquist, S. (2007) A suite of Gateway® cloning vectors for high-throughput genetic analysis in Saccharomyces cerevisiae. Yeast, 24, 913–919.

50. Lousa, M., Vázquez-Huarte-Mendicoa, C., Gutiérrez, A.J., Saavedra, P., Navarro, B. and Tugores, A. (2019) Genetic epidemiology, demographic, and clinical characteristics of Charcot-Marie-tooth disease in the island of Gran Canaria (Spain). Journal of the Peripheral Nervous System, 24, 131–138.

51. Sun, L., Song, Y., Blocquel, D., Yang, X.-L. and Schimmel, P. (2016) Two crystal structures reveal design for repurposing the C-Ala domain of human AlaRS. Proceedings of the National Academy of Sciences of the United States of America, 113, 14300–14305.

52. Karczewski, K.J., Francioli, L.C., Tiao, G., Cummings, B.B., Alföldi, J., Wang, Q., Collins, R.L., Laricchia, K.M., Ganna, A., Birnbaum, D.P., et al. (2020) The mutational constraint spectrum quantified from variation in 141, 456 humans. Nature, 581, 434–443.

53. Qin, X., Deng, X., Chen, L. and Xie, W. (2016) Crystal Structure of the Wild-Type Human GlyRS Bound with tRNA(Gly) in a Productive Conformation. Journal of Molecular Biology, 428, 3603–3614.

54. Koh, C.Y., Wetzel, A.B., de van der Schueren, W.J. and Hol, W.G.J. (2014) Comparison of histidine recognition in human and trypanosomatid histidyl-tRNA synthetases. Biochimie, 106, 111–120.

55. Austin, J. and First, E.A. (2002) Catalysis of tyrosyl-adenylate formation by the human tyrosyl-tRNA synthetase. J Biol Chem, 277, 14812–14820.

56. Yang, X.-L., Otero, F.J., Ewalt, K.L., Liu, J., Swairjo, M.A., Köhrer, C., RajBhandary, U.L., Skene, R.J., McRee, D.E. and Schimmel, P. (2006) Two conformations of a crystalline human tRNA synthetase-tRNA complex: implications for protein synthesis. EMBO J, 25, 2919–2929.

57. Raimondi, C., Brash, J.T., Fantin, A. and Ruhrberg, C. (2016) NRP1 function and targeting in neurovascular development and eye disease. Prog Retin Eye Res, 52, 64–83.

58. Brachmann, R.K., Vidal, M. and Boeke, J.D. (1996) Dominant-negative p53 mutations selected in yeast hit cancer hot spots. Proc Natl Acad Sci U S A, 93, 4091–4095.

59. Marutani, M., Tonoki, H., Tada, M., Takahashi, M., Kashiwazaki, H., Hida, Y., Hamada, J., Asaka, M. and Moriuchi, T. (1999) Dominant-Negative Mutations of the Tumor Suppressor p53 Relating to Early Onset of Glioblastoma Multiforme. Cancer Res, 59, 4765–4769.

60. Quimby, B.B., Alano, A., Almashanu, S., DeSandro, A.M., Cowan, T.M. and Fridovich-Keil, J.L. (1997) Characterization of two mutations associated with epimerase-deficiency galactosemia, by use of a yeast expression system for human UDP-galactose-4-epimerase. Am J Hum Genet, 61, 590–598.

61. Elsevier, J.P. and Fridovich-Keil, J.L. (1996) The Q188R mutation in human galactose-1-phosphate uridylyltransferase acts as a partial dominant negative. J Biol Chem, 271, 32002–32007.

