## Supplemental Materials for "A humanized yeast model reveals dominant-negative properties of neuropathy-associated alanyl-tRNA synthetase mutations"

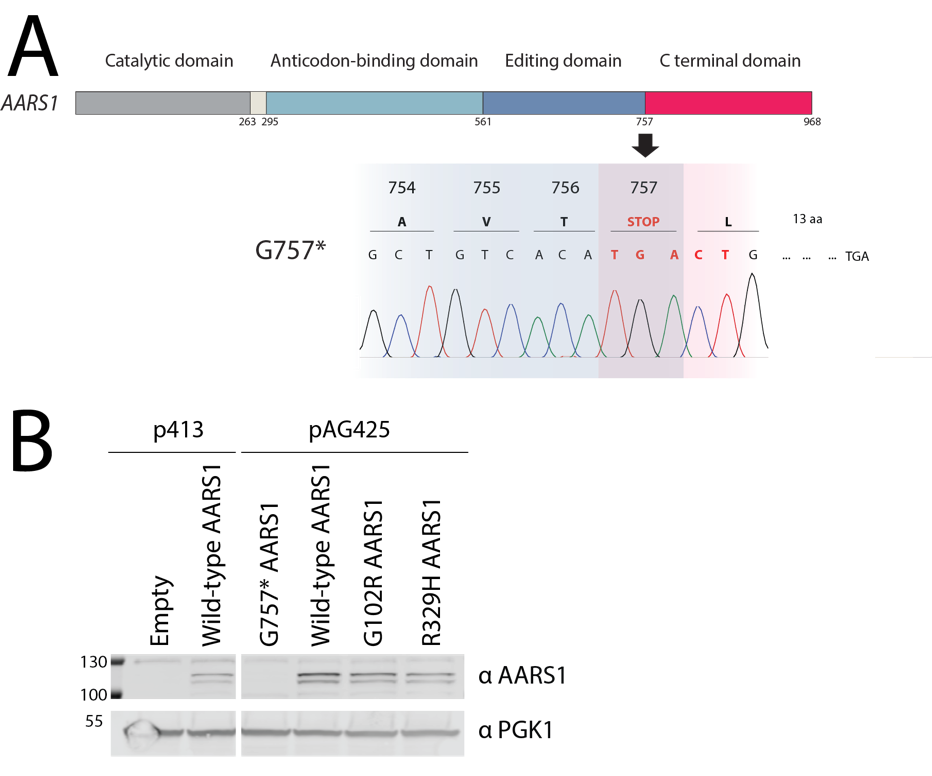

**Supplemental Figure 1.** Yeast expression of human wild-type or mutant AARS1. **(A)** A cartoon of the AARS1 protein structure highlights the junction between the editing domain (blue) and the C-terminal domain (pink). A chromatogram illustrates the five base pair insertion introduced at this junction to create a premature stop codon, G757*. This insertion also shifts the open reading frame to generate another stop codon 13 amino acids downstream. **(B)** Yeast were transformed with a p413 vector, either with no insert or expressing human wild-type *AARS1*, or were transformed with a pAG425 vector expressing wild-type or mutant human *AARS1*. Yeast were grown in galactose and raffinose media lacking either histidine or leucine, with no doxycycline. A western blot was performed with yeast lysates; the vectors present in each yeast strain are indicated across the top. Protein molecular weights are indicated along the left side in kilodaltons, and antibodies are indicated on the right side. A representative image from four biological replicates is shown. Of note, full-length AARS1 is predicted to migrate at 107kDa. The lower band likely corresponds to a downstream open-reading frame beginning at M44, which is predicted to produce a 102kDa AARS1 protein.

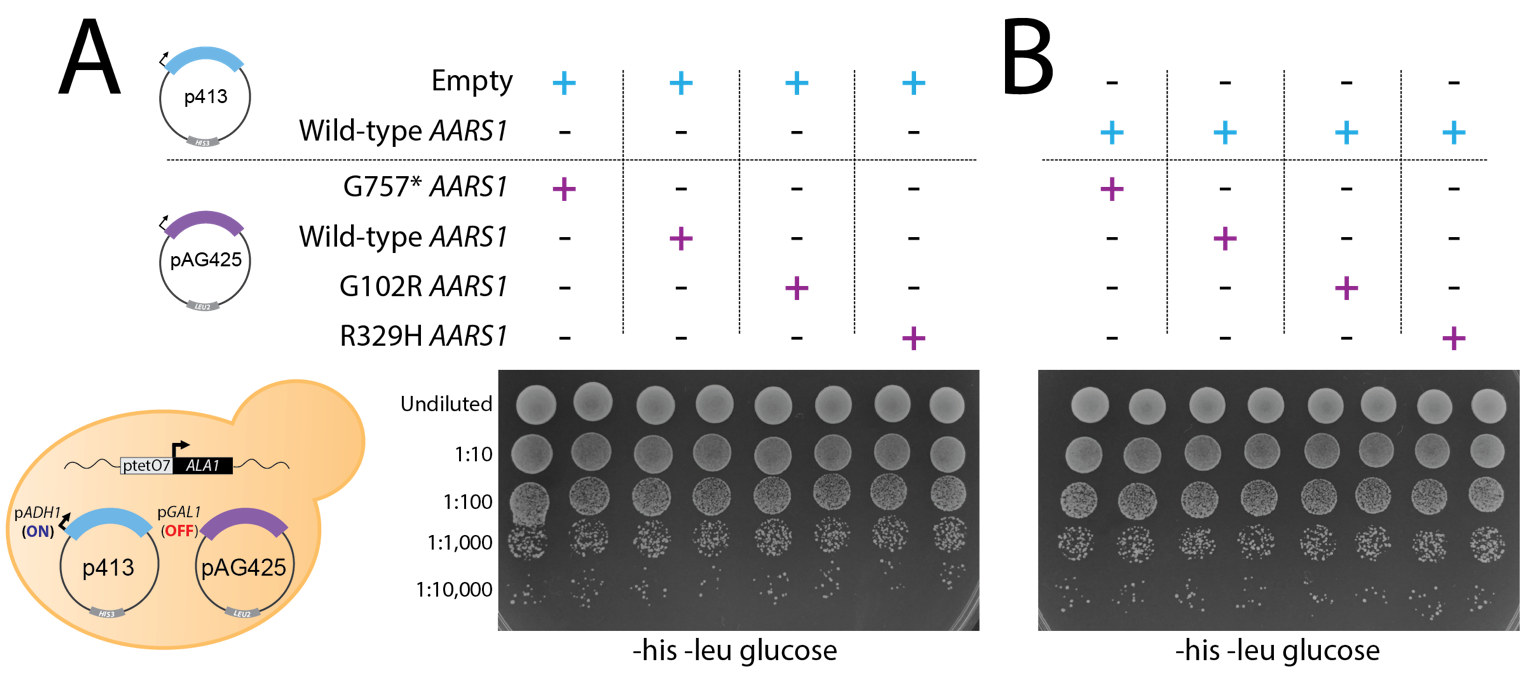

**Supplemental Figure 2.** Yeast expressing G102R and R329H *AARS1* do not exhibit growth defects on glucose media lacking doxycycline. (**A**) Yeast harboring an endogenous doxycycline-repressible *ALA1* locus were transformed with a p413 vector with no insert and a pAG425 vector to express either wild-type or mutant human *AARS1*. Cultures were plated undiluted or diluted on media lacking histidine and leucine, and containing glucose as the carbon source. (**B**) Similar to panel A, except that wild-type human *AARS1* is expressed from p413. For both panels, the vectors present in each experiment are indicated across the top, the dilution of the spotted yeast cultured is indicated on the left, and the media conditions are indicated across the bottom (his = histidine; leu = leucine). Representative images are shown from thirteen (for G102R) or sixteen (for R329H) biological replicates. A cartoon on the bottom left illustrates the experimental conditions for all samples.

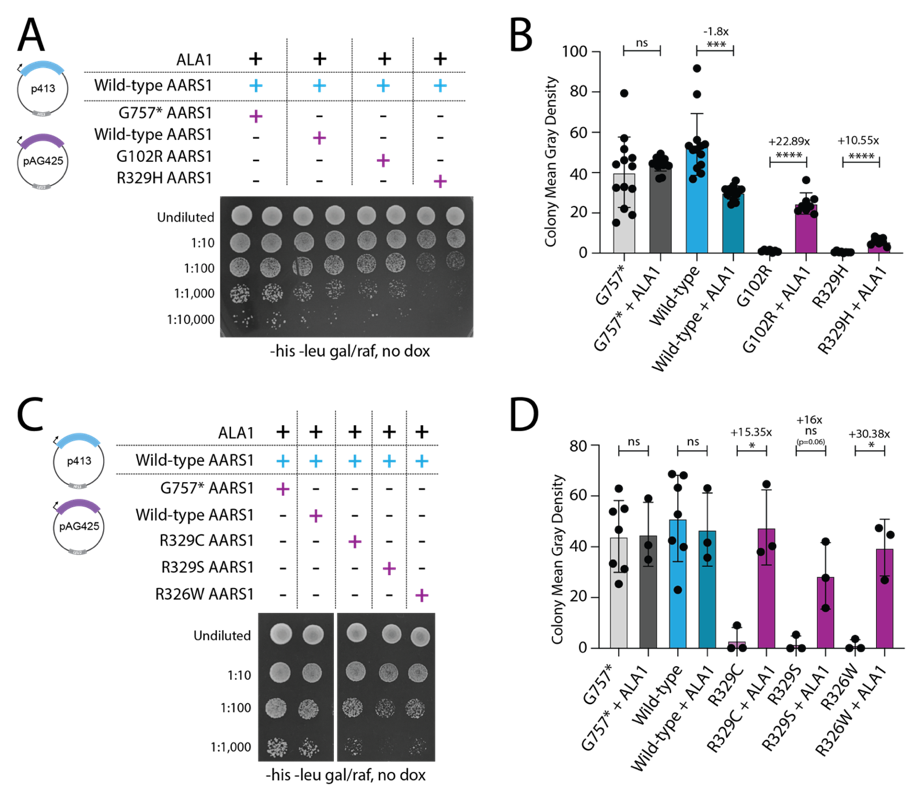

**Supplemental Figure 3.** *ALA1* expression rescues yeast growth. **(A** and **C**) Yeast with the doxycycline-repressible endogenous *ALA1* locus were transformed with a p413 vector expressing wild-type human *AARS1*, as well as a pAG425 vector to express either wild-type or mutant *AARS1.* Cultures were plated undiluted or diluted on media lacking histidine and leucine, with galactose/raffinose as the carbon source. No doxycycline was included in the media. Vectors present in each experiment are indicated across the top, the dilution of the spotted yeast cultured is indicated on the left, and media conditions are shown across the bottom (his = histidine; leu = leucine; gal = galactose; raf = raffinose). **(B)** Colony intensity for the strains shown in **(A)** was quantified using ImageJ. The mean and standard deviation for at least thirteen biological replicates is shown. Quantification of the yeast that were plated on doxycycline media (images presented in Figure 1) is included for comparison. **(D)** ImageJ was used to quantify colony intensity of the colonies shown in **(C).** The mean and standard deviation for three replicates is shown. The quantification of the yeast that were plated on doxycycline (shown in Figure 7) is included for comparison. For **(B)** and (**D)**, the fold change between the mean colony intensity without *ALA1* and the mean colony intensity with *ALA1* is shown. Statistical significance was determined with a Welch’s t-test. **** p<0.0001, *** p<0.001, * p<0.05.

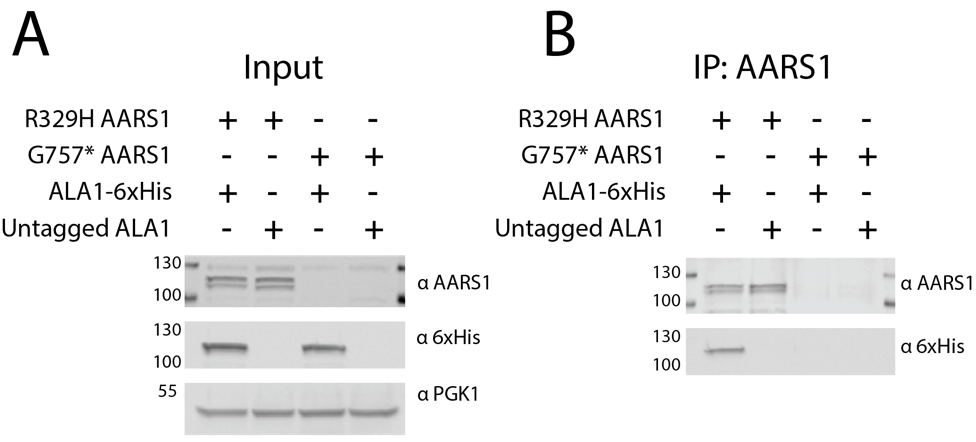

**Supplemental Figure 4.** R329H AARS1 interacts with ALA1. **(A)** Yeast were transformed with vectors to co-express R329H AARS1 and 6xHis-tagged ALA1. G757* AARS1 and untagged ALA1 were included as negative controls. A western blot was performed to detect the resulting proteins, along with the PGK1 loading control. The presence or absence of each construct is shown across the top of the image, the protein molecular weights are indicated in kilodaltons (kDa) across the left of the image, and the antibodies are shown to the right of the image. **(B)** After immunoprecipitation with an AARS1 antibody, a western blot was performed to detect co-immunoprecipitated protein. The image is annotated as in panel A. A representative image from two technical replicates is shown.

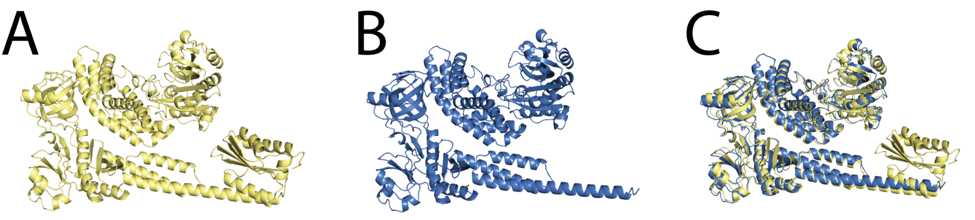

**Supplemental Figure 5.** Q855* *AARS1* does not significantly change the predicted AARS1 protein structure outside of the C-terminal globular domain. **(A)** The predicted structure of full-length AARS1, determined using AlphaFold Colab, is depicted in yellow. **(B)** The predicted structure of Q855* AARS1, determined using AlphaFold Colab, is depicted in blue. **(C)** Predicted structures from **(A)** and **(B)** are superimposed.

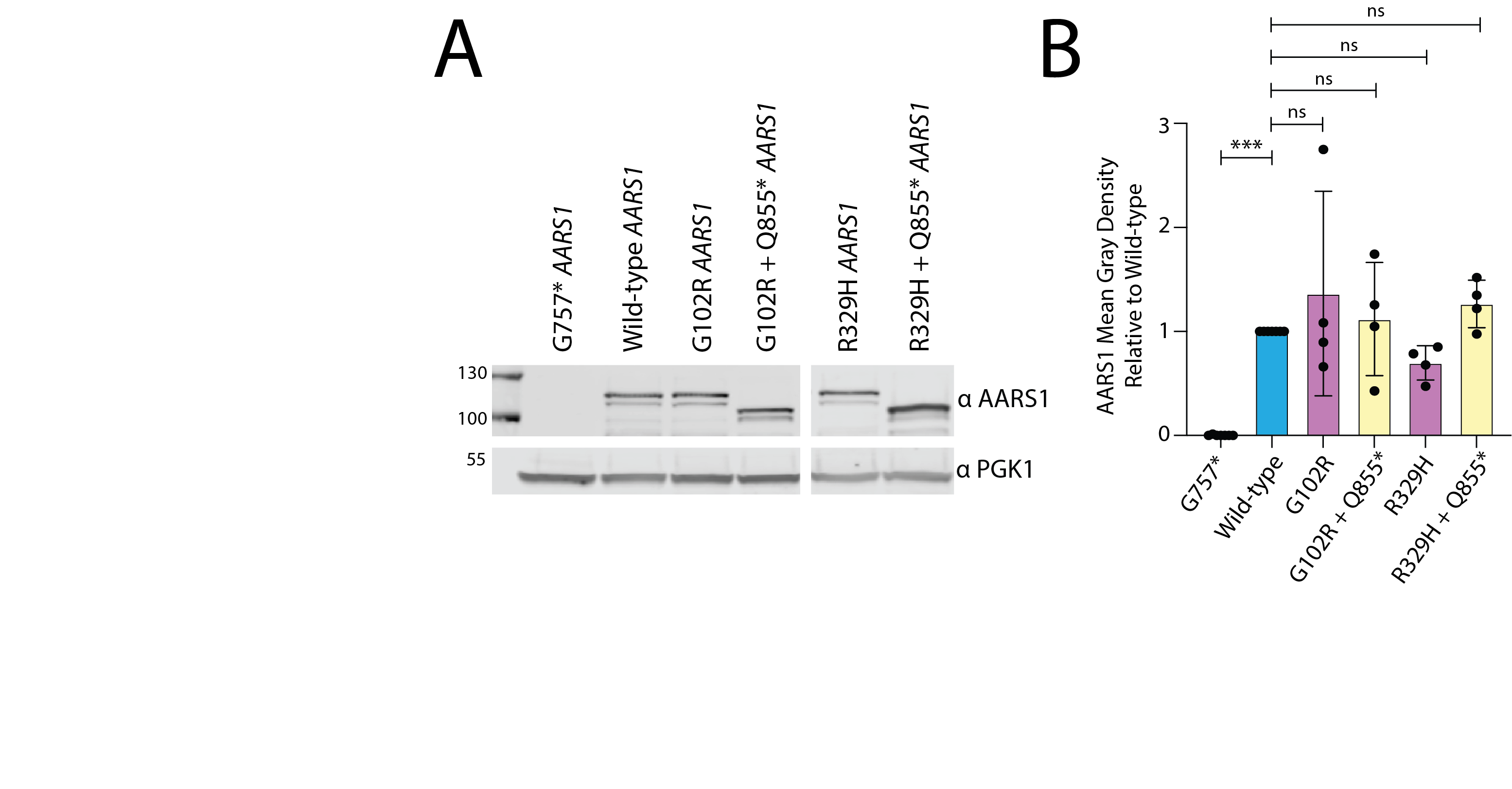
**Supplemental Figure 6.** Q855* *in cis* with G102R or R329H does not impair AARS stability. **(A)** Yeast harboring a doxycycline-repressible endogenous *ALA1* locus were transformed with a pAG425 vector to express either wild-type or mutant human *AARS1*. Yeast were grown in galactose and raffinose media lacking leucine, with no doxycycline. Protein lysates were subjected to western blot analysis to detect the resulting human AARS1 proteins. The constructs present in each experiment are indicated across the top of the image, protein molecular weight is indicated in kilodaltons on the left of the image, and antibodies are indicated on the right of the image. **(B)** ImageJ quantification of western blots. Bars indicate the mean and one standard deviation from three biological replicates. A one-way ANOVA with Dunnett’s multiple comparisons test was performed to determine if protein levels of each AARS1 variant significantly differed from protein levels of wild-type AARS1*.*

| Patient ID | Gender | Age | Mutation | Age of Onset | CMTNS | Vibration LL | Vibration UL | Cutaneous LL | Cutaneous UL | Ulnar DML (ms) <3.4 | Ulnar NCV1 (m/s) >49 | Ulnar NCV2 (m/s) >50 | Ulnar CMAP (mV) >2.8 | Median DML (ms) <3.5 | Median NCV (m/s) >48 | Median CMAP (mV) >3.5 |
| --- | --- | --- | --- | --- | --- | --- | --- | --- | --- | --- | --- | --- | --- | --- | --- | --- |
| 75872-0001 | Male | 56 | R329S | 11 | 29 (Severe) | Absent toes, ankles, knees | Absent fingers, reduced wrists and elbows | Absent toes, ankles, knees | Absent fingers, wrists | 4.4 | 24 | 23 | 0.323 | NR | NR | NR |
| 75292-0001 | Male | 76 | R329H | 56 | 11 (Low moderate) | Reduced toes, ankles, knees | Normal | Normal | Normal | 3.2 | 39 | 42 | 9.3 | 5.4 | 39 | 5.4 |
| 75292-9001 | Female | 44 | R329H | 20 | 7 (Mild) | Reduced toes, ankles, knees | Normal | Normal | Normal | 2.9 | 49 | 50 | 8 | 4.5 | 43 | 4.9 |
| 75292-0100 | Male | 72 | R329H | 25 | 11 (Low moderate) | Absent toes, reduced ankles | Normal | Normal | Normal | 3.2 | 45 | 55 | 10.1 | 4.8 | 41 | 5.6 |

**Supplemental Table 1.** Clinical presentation of four individuals with *AARS1*-mediated dominant peripheral neuropathy. CMTNS=CMT Neuropathy Score, LL=lower limb, UL=upper limb, DML=distal motor latency (upper limit of normal is 3.4 milliseconds for the ulnar nerve and 3.5 milliseconds for the median nerve), NCV=nerve conduction velocity (lower limit of normal is 49 or 50 meters per second for the ulnar nerve and 48 meters per second for the median nerve), CMAP = compound muscle action potential (lower limit of normal is 2.8 millivolts for the ulnar nerve and 3.5 for the median nerve). NR=no recording
